# Programmable nanobody circuits for cell selection

**DOI:** 10.64898/2026.02.22.707289

**Authors:** Nathan B. Wang, Albert Blanch-Asensio, Hannah Cevasco, Deon S. Ploessl, Derin R. Gumustop, Mary E. Ehmann, Maria F. Castellanos, Francisco J. Sánchez-Rivera, Timothy M. O’Shea, Kate E. Galloway

**Author notes:** Correspondence (K.E.G.).

## Abstract

Efficient and scalable isolation of specific cell populations remains a central bottleneck for genome engineering, pooled screening, and cell therapy manufacturing. Here, we present DASIT (**<u>D</u>**estabilized-nanobody **<u>A</u>**ntigen **<u>S</u>**election and **I**dentification **<u>T</u>**ool), a protein-based circuit for antigen-specific cell selection. DASIT uses a destabilized nanobody fused to an antibiotic resistance protein. In cells expressing the target antigen, binding of the nanobody fusion to the cognate antigen stabilizes DASIT, thereby coupling the presence of an antigen to a selectable signal. We developed DASIT circuits that enable robust selection of antigen-expressing cells and show that they can be designed to target distinct antigen classes and perform across cell types. Because DASIT operates at the protein level, it supports both stable integration and transient delivery, enabling recyclable selection without permanent genomic integration of resistance markers. We demonstrate scalable, FACS-free enrichment in three challenging applications: multiplexed, logic-gated integration of landing pads in human iPSCs, high-throughput CRISPR screening, and phenotypic selection of *in vitro*-derived neurons at transplantation scale. By decoupling selection from vector integration, DASIT establishes an automation-compatible architecture for multistep genome engineering, high-throughput library screening, and large-scale cell manufacturing.

**Highlights:**

DASIT enriches for antigen-positive cells across multiple selection markers and antigens
Intermediate levels of DASIT expression support selection across stable and transient delivery modalities
Logic-gated, precision genome engineering of human iPSCs via DASIT selection
DASIT enables scalable activity-based selection for high-throughput base editing screens
DASIT-selected engineered motor neurons survive grafting into acute spinal cord injury

---

Selection and isolation are critical steps in the production of cells for phenotypic screening, drug screening, bioproduction, and cell therapies. The explosion of precision DNA-editing tools and engineering approaches for producing “designer” mammalian cells has the potential to transform biomedicine^1,2^. Cells can be edited *in vitro* for disease modeling, or even *in vivo* to directly treat diseases^1^. In addition, cells can be isolated or generated *in vitro*, edited, and transplanted into patients for cell-based therapies^3–7^. However, each of these processes results in mixed populations. While engineered cells have extraordinary potential for biomedicine, there is a critical need for scalable methods of cell isolation of desired cell types, genotypes, and phenotypes. Fluorescence-activated cell sorting (FACS) and magnetic-activated cell sorting (MACS) can enrich specific cells based on the presence (or absence) of a detectable signal. However, the harsh mechanical forces of these flow-based methods present a major drawback that can lead to substantial cell loss and reduced viability for fragile cell types^8,9^. Further, these sorting processes are inherently limited by scale. Time and selection resources increase as the total number of cells increases, becoming prohibitive at the large scales required for massive libraries and cell therapies^10–12^. The lack of scalable methods for isolation of *in vitro*-engineered cells significantly increases costs and limits their broader use^6^. Antibiotic selection offers a scalable method of selecting for transgenic cells but generally requires the integration of selection markers which can limit downstream use of these selections. Given the small set of effective selection markers, marker recycling is essential for iterative genome writing workflows^13–16^.

To overcome the limitations of traditional selection markers, we identified destabilized nanobodies (desNb)—also referred to as conditionally stable nanobodies^17^—as candidate molecules that could select for cells based on expression of a specific antigen. DesNbs are small single-domain antibodies that are stabilized when bound to their antigen and degraded in the absence of antigen^17^. DesNbs can be generated from wildtype nanobodies by introducing destabilizing mutations in conserved framework regions^17^. By fusing desNbs to effector proteins, cognate antigen inputs can be transformed into specific functions defined by the effector protein, such resistance conferred by a selection enzyme (Fig 1A)^17^. As desNb and effector domains are modular, each module can be optimized to tune specificity and activity. Importantly, because desNbs function at the protein level, these protein-based circuits are compatible with diverse delivery methods and can rapidly respond to antigen levels. Existing protein-based circuits largely rely on proteases, degrons, and synthetic protein interactions to compute responses^18–25^. Despite their potential as sensors, nanobodies have seen limited use in protein circuits^17,25–30^.

**Figure 1.**
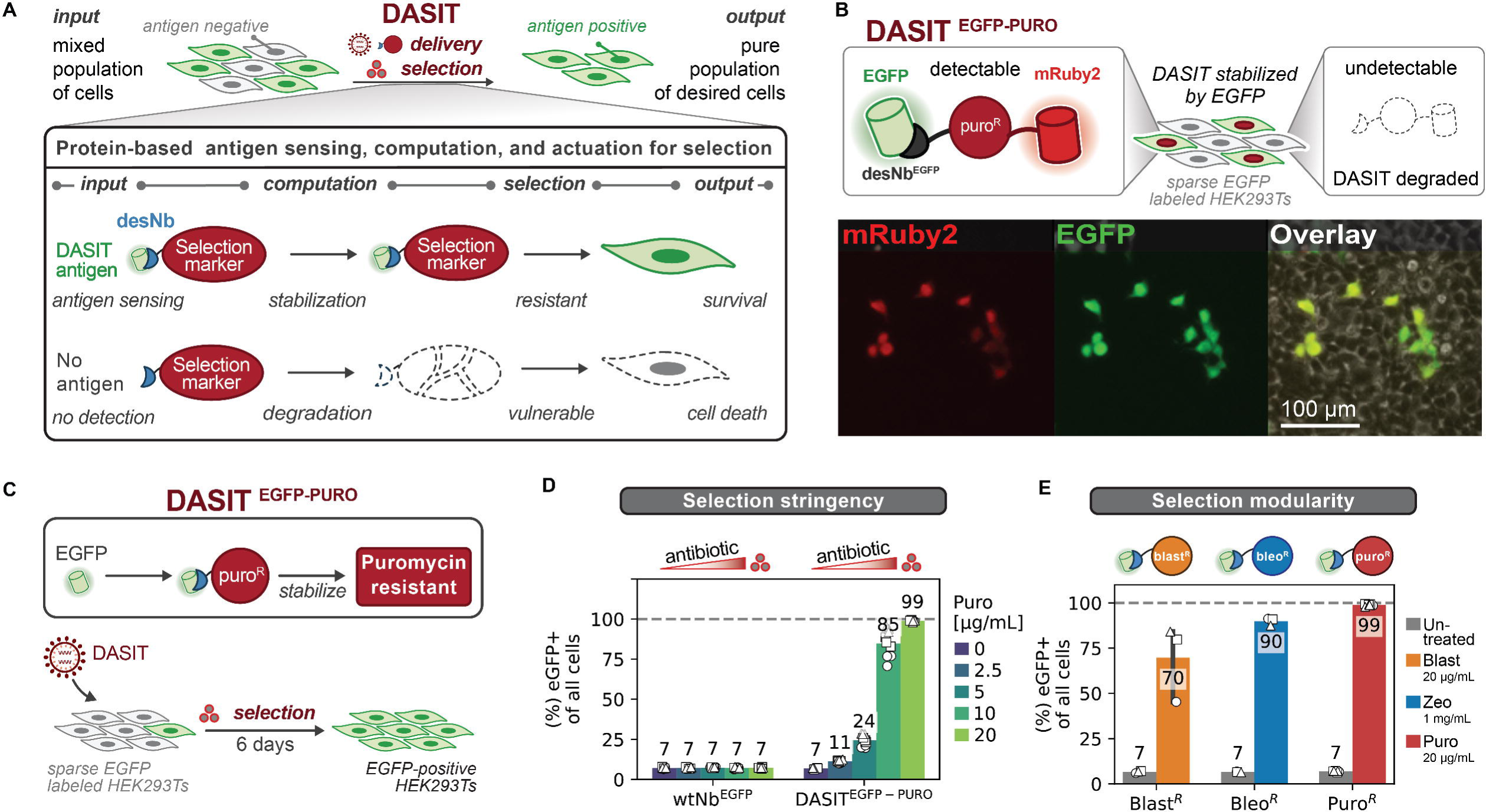
DASIT enriches for antigen-positive cells across multiple antibiotic resistance markers and antigens. A. Destabilized nanobodies (desNb’s) convert protein-level antigen inputs into specific output actions by fusion to various effectors. DASIT fuses desNb’s to antibiotic resistance proteins to convert cells with desired intracellular antigens into a selectable signal. B. Images of sparsely labeled EGFP-positive and EGFP-negative HEK293T cells transduced with DASIT^EGFP-PURO^ (desNb^EGFP^-puroR-mRuby2) is only stabilized in EGFP+ cells. Images of mRuby2 (left), EGFP (middle), and overlaid with phase contrast (right) are shown. C-D. (C) DASIT^EGFP-PURO^ delivery into sparsely EGFP labeled HEK293T cells with puromycin selection for 6 days. (D) EGFP purity after selection with different puromycin concentrations. Mean is shown and markers denote biological replicates; n ≥ 3 biological replicates per condition; dashed line marks 100% purity. E. DASIT delivery at MOI=5 into single-EGFP integrated HEK293T for 3 different resistances: blasticidin, zeocin, and puromycin. EGFP+ purity with and without antibiotic treatment was quantified after 6 days of puromycin selection and 10 days of blasticidin and zeocin selection (see Fig S1C for 6 vs 10 day purities). Mean is shown and markers denote biological replicates; n ≥ 3 biological replicates per condition; dashed line marks 100% purity.

In this work, we developed a novel method for scalable selection of engineered cells called **<u>D</u>**estabilized-nanobody **<u>A</u>**ntigen **<u>S</u>**election and **I**dentification **<u>T</u>**ool (DASIT). DASIT is a protein-level circuit that uses desNbs that bind to an intracellular antigen specifically expressed in desired cells. By fusing an antibiotic resistance gene to the desNb, we confer selective resistance to cells that express the desired antigen. Both stable and transient delivery of DASIT supports selection in diverse cells including human induced pluripotent stem cells (iPSCs) and cancer cell lines, supporting rapid logic-gated genome engineering in iPSCs and functional enrichment for CRISPR base-editing activity to facilitate massive library screening. Finally, we demonstrate that DASIT can select for cell identity, allowing us to purify *in vitro*-engineered motor neurons from a mixed culture of reprogrammed cells and graft these cells following acute spinal cord injury in mice. Given its modularity, we anticipate that the DASIT framework is extensible to diverse applications from cell engineering to activity-based screening.

## RESULTS

### DASIT enriches for antigen-positive cells across multiple antibiotic resistance markers and antigens

To establish DASIT as a general strategy for antigen-dependent cell enrichment, we first tested whether intracellular antigen binding could be converted into selective cell survival. Since DASIT relies on conditional stabilization of an antibiotic resistance protein, effective selection requires the fusion protein to be specifically stabilized in antigen-positive cells while remaining degraded in antigen-negative cells. We therefore evaluated DASIT in a mixed control population of cells expressing or lacking EGFP to assess antigen-specific stabilization and enrichment. We constructed DASIT^EGFP-PURO^ circuits using a well-established desNb that binds to EGFP (desNb^EGFP^). To confirm EGFP-specific stabilization, we transduced vectors expressing the desNb^EGFP^ fused to puro^R^-mRuby2 (desNb^EGFP^- puro^R^ -mRuby2) into a HEK293T line that was sparsely labeled with EGFP. As expected, mRuby2 was stabilized in only EGFP-positive cells, demonstrating that we could use the desNb framework to identify antigen-positive cells as previously demonstrated (Fig 1B)^17^.

With the ability to identify antigen-positive cells, we hypothesized that fusion of the antibiotic resistance marker to the destabilized nanobody could confer selective resistance to cells expressing EGFP. To test enrichment of EGFP-positive cells, we transduced DASIT^EGFP-PURO^ into a mixed population of non-transgenic and single-copy, EGFP-positive HEK293T cells and applied puromycin selection for 6 days (Fig 1C, S1A). As controls, we included a wildtype nanobody for EGFP (wtNb^EGFP^-puro^R^) which is stable in the absence of EGFP. As expected, delivery of the wtNb control did not enrich for EGFP-positive cells (Fig 1D). In contrast, selection with DASIT^EGFP-PURO^ shows a dose-dependent enrichment in EGFP-positive cells, generating a 14-fold enrichment and near complete selectivity (∼99%) at the highest amount of puromycin tested (Fig 1D). While this selection increased the fraction of EGFP-positive cells, we did not observe a significant change in the total number of cells at 6 days post treatment (dpt), suggesting that growth of EGFP-positive cells compensated for loss of EGFP-negative cells (Fig S1B). Next, we tried different antibiotic resistance proteins to test whether DASIT is compatible with different resistance mechanisms. Blast^R^ confers resistance to blasticidin and bleo^R^ confers resistance to zeocin, a member of the bleomycin family. As expected, both DASIT^EGFP-BLAST^ and DASIT^EGFP-BLEO^ enriched for EGFP-positive cells, but these circuits showed lower selectivity compared to DASIT^EGFP-PURO^ at 6 dpt (Fig 1E, S1C). In HEK293Ts, we could increase selectivity of DASIT^EGFP-BLAST^ and DASIT^EGFP-BLEO^ by extending the selection to 10 days (Fig S1C). Extended selection increases EGFP-positive purity up to ∼70% and ∼90% for blasticidin and zeocin, respectively, at 10 dpt (Fig 1E). These results demonstrate that DASIT is modular across different resistance markers.

In theory, desNbs can be rapidly generated from existing nanobodies by introducing destabilizing mutations in conserved framework regions^17^. Three major destabilization mutations were previously reported to be broadly applicable to several nanobodies, and three additional mutations—resulting in six total—increased specificity for identifying cells expressing EGFP^17^. We hypothesized that other nanobodies could be compatible with DASIT by transfer of destabilizing mutations. We first tested whether DASIT could respond to the presence of an epitope tag. Epitope tags are small, enabling detection, manipulation, and purification of exogenous and edited endogenous proteins. For proof of principle, we used the rationally designed, minimal, 13-residue ALFA-tag and its corresponding nanobody, which have previously been used for live-cell identification (Fig S2A)^31,32^. To generate a destabilized nanobody, we applied the three major mutations into the framework regions of the ALFA-tag nanobody (Fig S2B). We generated a population of HEK293T cells sparsely labeled with a single copy of a fluorescent TagBFP^ALFA^ reporter via lentiviral transduction. We transduced these cells with DASIT^PURO-ALFA^ along with a stable iRFP670 transduction marker (Nb^ALFA^-puro^R^-mRuby2-2A-iRFP670) to measure fluorescent protein stabilization. We were surprised to observe that the wtNb^ALFA^ showed selective fluorescent stabilization to TagBFP^ALFA^ while desNb^ALFA-3maj^ did not (Fig S2C).

To test another nanobody-antigen pair, we transferred the predicted destabilizing mutations onto a nanobody for mCherry (LaM4, i.e., wtNb^mCh^)^33^ (Fig S2D-E). Again, we measured fluorescent protein stabilization in a mixed population, first transducing HEK293T cells with mCherry and then transducing several DASIT variants targeting mCherry. Across all nanobody variants (wtNb^mCh^, desNb^mCh-3maj^, desNb^mCh-5mut^, desNb^mCh-6mut^), we did not observe mCherry-specific stabilization (Fig S2F). Altogether, we find that a simple mutation transfer does not predictably generate nanobody fusion proteins that are conditionally stable in the presence of their cognate antigen. Instead, the stability of nanobody fusions must be tested empirically. The exact properties of the nanobody-antigen interaction, the nanobody fusion protein, and the antibiotic resistance marker likely influence the efficacy of selection in the DASIT framework. Based on the fluorescent protein stabilization data, we concluded that wtNb^ALFA^ could be used for DASIT^ALFA^ selection.

### Intermediate levels of DASIT expression support selection for stable and transient delivery modalities

Given that DASIT functions at the protein level, we wondered whether we could transiently deliver DASIT circuits and enrich for engineered cells. A major benefit of this strategy is that selection markers can be recycled for subsequent engineering steps once the transiently delivered DASIT components are degraded. However, transient delivery may not support optimal expression of DASIT to enable selection. To ensure selectivity, off-target expression of the antibiotic resistance marker must be low. To confer resistance, expression of the resistance marker must be sufficient. Thus, selective resistance likely requires well-tuned expression of DASIT.

To compare transient delivery methods to our previous lentiviral-based delivery, we built a single transcript encoding DASIT^EGFP-PURO^ fused to a TagBFP output followed by a stable iRFP670 fluorescent reporter (Fig 2A). We used the iRFP670 fluorescent reporter to assess expression and delivery of the transcript via lentivirus, plasmid transfection, and modRNA transfection. As controls, we included the wildtype nanobody for EGFP (wtNb^EGFP^-puro^R^-TagBFP) which is stable in the absence of EGFP (Fig S3A-B). Using flow cytometry, we quantified the fluorescent selectivity of DASIT^EGFP-PURO^ by measuring how many cells are both EGFP-positive and TagBFP-positive normalized by the number of TagBFP-positive cells after gating for delivery (Fig 2B). As plasmid delivery results in the highest expression of the iRFP670 marker (Fig 2C, middle,, S3C), we hypothesize that high expression of DASIT due to high copy number from DNA transfection, leads to saturation of the cells’ degradation machinery^34^, preventing degradation of unbound DASIT^EGFP-PURO^. Conversely, when DASIT was delivered via lentivirus (MOI=5) or by modRNA transfection, expression of the co-delivery marker is lower and the fluorescent selectivity for desNb^EGFP^ increases to ∼97% and ∼95%, respectively. Thus, we find that there exists an optimal window of expression for DASIT^EGFP-PURO^ to support selection, one that is high enough to unlock the function of the fused output protein but low enough to avoid antigen-independent activity (Fig 2B-D).

**Figure 2.**
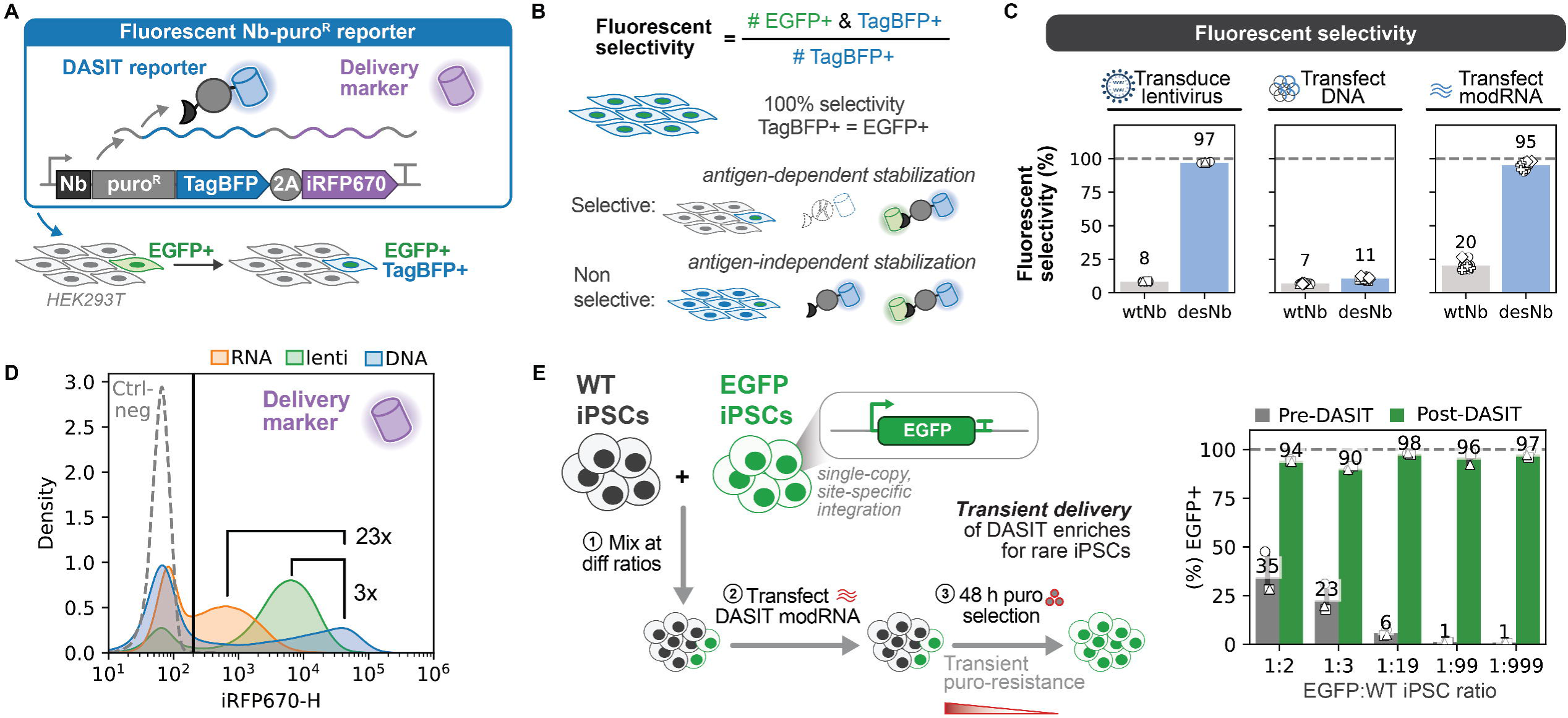
Intermediate levels of DASIT expression support selection for stable and transient delivery modalities. A. TagBFP fluorescence enables live identification of DASIT expression and a 2A-iRFP670 marker can be used as a co-expression marker. Single EGFP-integrated HEK293T cells support testing of DASIT selectivity based on EGFP and TagBFP fluorescence. B. Fluorescent selectivity is defined as the proportion of EGFP- and TagBFP-positive cells of the TagBFP-positive cells. desNb^EGFP^-puro^R^-TagBFP should only stabilize and express in EGFP+ cells. Conversely, the non-selective wildtype (wtNb) control will be stable in EGFP-negative and EGFP-positive cells. C. DASIT fluorescent selectivity after gating delivered cells using the stable iRFP670 marker. The fluorescent selectivity was measured for single EGFP-integrated 293Ts (∼7% EGFP+) after delivery by lentiviral transduction (left), DNA transfection (middle), and RNA transfection (right), and for wtNb^EGFP^ and desNb^EGFP^. Mean is shown with 95% confidence interval and markers denote biological replicates; n ≥ 3 biological replicates per condition; dashed line marks 100% purity. D. Representative iRF670 fluorescence distribution (see Fig S3C for replicates). DNA was measured at 48 hpt, RNA at 24 hpt, and lentivirus >6 dpi. Dashed line indicates the negative control fluorescence. E. Schematic of transient enrichment of rare cell populations using DASIT (left) and flow cytometry analysis before and after DASIT selection of different cell populations at varying ratios of EGFP+ to negative cells. Mean is shown with 95% confidence interval and markers denote biological replicates; n = 3 biological replicates per condition; dashed line marks 100% purity.

To determine whether transient DASIT expression could enable rapid enrichment of rare, engineered cells, we tested whether modRNA delivery could selectively enrich human iPSCs carrying a single, site-specifically integrated copy of a transgene. Genetically engineered iPSCs are useful in many biomedical applications and enrich rapidly (within 48 hours) using certain antibiotic resistance markers^35,36^, making them an ideal application to test transient DASIT selection. To do so, we mixed populations of non-transgenic (wt) iPSCs and transgenic iPSCs with a single copy of EGFP at the *CLYBL* safe harbor locus at ratios ranging from 1:2 to 1:999 of EGFP:wt cells (Fig 2E). We delivered DASIT^EGFP-PURO^ as modRNA to these mixed populations and applied selection for 48 hours. Across all tested ratios, selection increases the EGFP-positive purity of the mixed iPSC cultures to ≥90%. Thus, RNA delivery of DASIT can enrich rare populations of engineered iPSCs, paving the way for accelerating cell engineering workflows through transient, rapid selection.

### Logic-gated, precision genome engineering of human iPSCs via DASIT selection

Having established that we could enrich rare populations of engineered human iPSCs, we wondered if DASIT could accelerate site-specific genome engineering. As a demonstration, we chose installation of a landing pad system^36^. Landing pad systems rapidly integrate large DNA payloads at pre-installed attP recombination sites using large serine recombinases^35,37,38^. While landing pad cell lines are extremely useful, establishing these lines requires months of engineering and validation^35,36,39^. Cargo integration at landing pads, such as the STRAIGHT-IN system, achieves high efficiency by selection via a promoter trap on the donor DNA payload that induces expression of a selection marker once cells integrate the DNA payload^35,36,39^. Use of these markers to select landing pad cells for DNA payload integration means those same markers cannot be used for the initial genome engineering step for landing pad integration, a challenge for iterative cell engineering workflows including large-scale genome rewriting^13–16^. Instead of selection, clones integrated with landing pads are generally isolated by FACS based on expression of a fluorescent reporter encoded in each landing pad.^39^ However, sorting increases cell stress, contamination risks, and recovery timelines which are intensified by the single-cell cloning step^39^. To circumvent this issue, we hypothesized that we could use transient DASIT enrichment to rapidly establish landing pad lines in novel donor iPSC lines.

As proof of principle, we set out to generate novel landing pad iPSC lines in a new human donor iPSC line. We chose iPS11, a commonly used human iPSC line^38,40–46^. First, we confirmed that transient modRNA expression of DASIT was sufficient to enrich a single landing pad containing the GT attP site for Bxb1 and expressing a fluorescent protein that stabilizes desNb^EGFP^ (Fig S4A). We integrated the landing pad via TALEN-mediated *CLYBL* targeting^47^. Nine of ten clones screened had a single landing pad correctly integrated at *CLYBL* as shown by junction PCR and by ddPCR (Fig S4B-C). After selecting and expanding one clone, we confirmed that this monoclonal landing pad line supports single-copy integration and expression of fluorescent proteins and functional transcription factor cargoes at the landing pad for rapid cell-fate programming (Fig S5).

Given the capacity to integrate cargoes into either *CLYBL* allele, we next tested DASIT’s ability to perform rapid logic-gated genome engineering. By using two orthogonal attP sites (GT attP or GA attP) within landing pads at either *CLYBL* allele, the STRAIGHT-IN Dual system supports simultaneous integration of two independent, large, DNA payloads^36^. Previously, iPSC lines engineered using the STRAIGHT-IN Dual system were isolated by sequential integration and lengthy, weeks-long single-cell cloning steps. To more rapidly integrate the Dual system, we tested enrichment of human iPSCs that successfully integrated co-delivered STRAIGHT-IN Dual landing pads by transient DASIT delivery of two orthogonal DASIT systems, DASIT^EGFP-BLEO^ and DASIT^ALFA-PURO^ (Fig 3A, S6).

**Figure 3.**
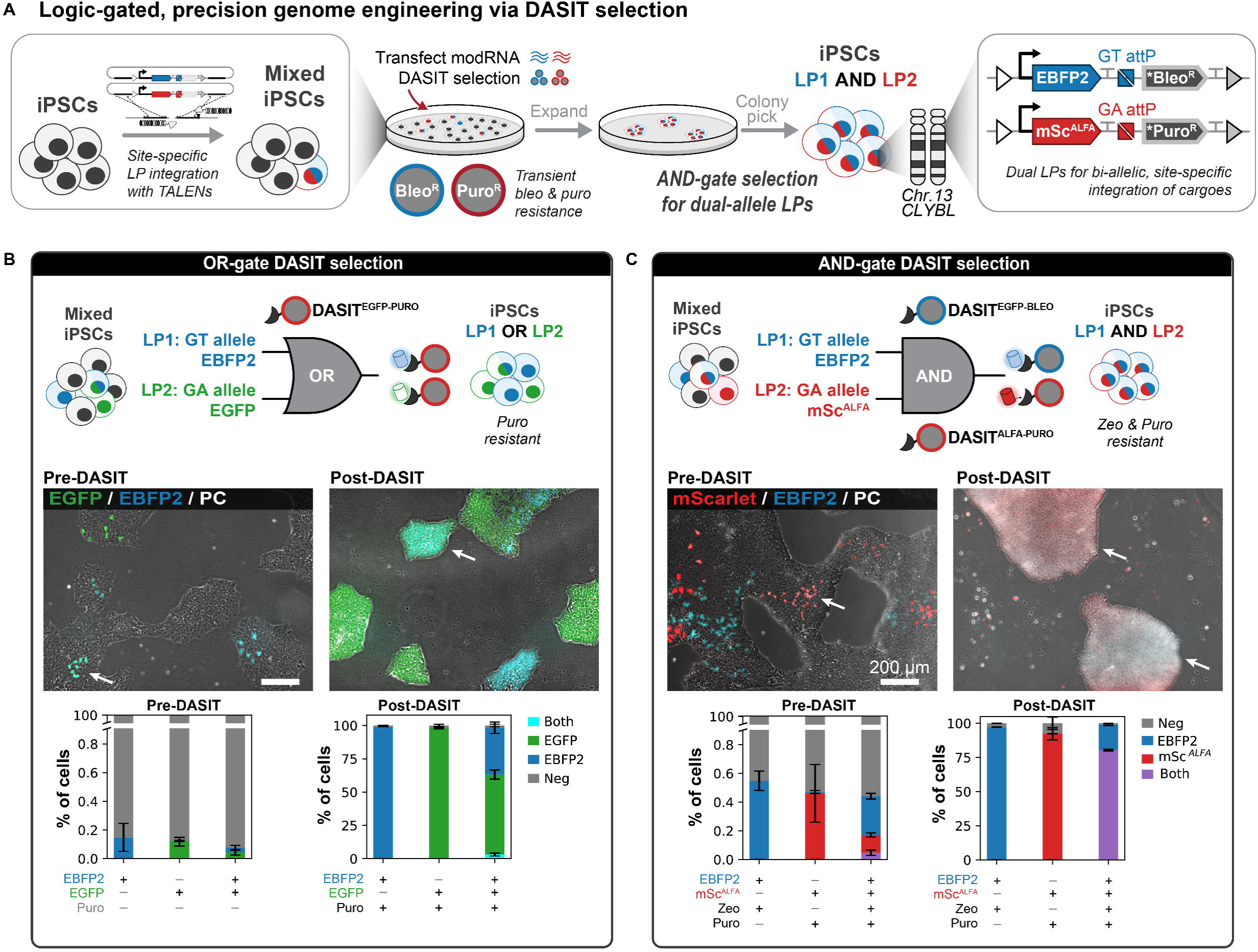
Logic-gated, precision genome engineering via DASIT selection. A. Schematic of how transient DASIT delivery and selection can be used to rapidly install and isolate iPSC clones with STRAIGHT-IN Dual landing pads. See Fig S6 for more details. B. Schematic of OR-gate strategy using DASIT selection to install one to two landing pads in the same genome (*top*). Overlay of fluorescence and phase contrast images (*middle*) and flow cytometry analysis (*bottom*) showing EBPF2 and EGFP reporter expression before and after DASIT OR-selection. C. Schematic of AND-gate strategy using DASIT selection to install two landing pads in the same genome (*top*). Overlay of fluorescence and phase contrast images (*middle*) and flow cytometry analysis (*bottom*) showing EBPF2 and mScarlet^ALFA^ (mSc^ALFA^) reporter expression before and after DASIT AND-selection.

To select for landing pad integrated iPSCs, we explored two logic-based (OR, AND) selection strategies (Fig 3B-C). For OR logic, we attempted dual landing pad integration by co-delivery of landing pads that express either an EBFP2 reporter for the GT allele or an EGFP reporter for the GA allele (Fig 3B, top). Due to their structural similarities, DASIT^EGFP-PURO^ can be stabilized by EGFP-related fluorescent proteins such as EBFP2. Thus, DASIT^EGFP-PURO^ can select for cells that express EBFP2, EGFP, or both. As expected, modRNA transfection of DASIT^EGFP-PURO^ and selection with puromycin results in a mix of cells expressing EBFP2, EGFP, and both (Fig 3B, S7A-B). For AND logic, we co-delivered landing pads expressing an EBFP2 reporter for the GT allele and an mScarlet with a C-terminal ALFA-tag (mScarlet^ALFA^) reporter for the GA allele (Fig 3C, top). DASIT^ALFA-PURO^ selects for cells that have integrated the GA allele and express mScarlet^ALFA^. As expected, modRNA co-transfection of DASIT^EGFP-BLEO^ and DASIT^ALFA-PURO^ and co-selection with zeocin and puromycin enriches for cells that have integrated landing pads at both alleles (Fig 3C, S7C-D). To verify function of the landing pads in the DASIT-engineered dual line, we selected one iPSC clone to expand. Using this monoclonal iPS11 dual landing pad line, we delivered constitutively expressed mGreenLantern and iRFP670 fluorescent reporters to the GT and GA alleles, respectively (Fig S8A). Following selection, both DNA payloads expressed in ∼98% of the cells (Fig S8B-C). Thus, transient expression of DASIT followed by selection can rapidly isolate functional dual landing pad lines in human iPSCs. By avoiding FACS and supporting marker recycling, DASIT facilitates rapid, multi-step cell engineering in an architecture that transforms a time- and labor-intensive process into a more rapid, self-contained, and potentially automatable workflow.

### DASIT enables scalable activity-based selection for high-throughput base editing screens

Given DASIT’s performance in selecting rare, engineered iPSCs, we hypothesized that DASIT could enable scalable activity-based selection from a heterogenous population in a high-throughput genome editing screen. Base editing screens use complex single guide RNA (sgRNA) libraries to install disease-relevant single-nucleotide transition mutations for phenotypic interrogation^48^. Screen scale-up is challenging in part due to the inability to rapidly, efficiently, and iteratively select cells with robust genome editing activity. For example, if only 1% of transduced cells exhibit editing activity, the empirical library representation will be lower than expected by several orders of magnitude, and 99% of the cells will only contribute background noise, potentially confounding the genotype-to-phenotype mapping. To optimize this signal-to-noise ratio, CRISPR-based screens often rely on fluorescent markers or other constructs to enrich for cells that express library sgRNAs and have detectable editing activity^49,50^. Isolating this population, however, can be both challenging and time-consuming. A single screen replicate of a library composed of 15,000 guides can require >20 hours of continuous FACS to purify enough cells with detectable base editing activity to achieve a representation of ≥10,000x as required for a robust *in vivo* screen.

We asked whether DASIT-mediated selection could enrich for cells with active base editing while preserving the representation of a complex sgRNA library. To answer this, we conducted an *in vitro* adenine base editing (ABE) screen using a reporter that expresses EGFP protein upon successful editing at the EGFP start codon.^50^ We first stably integrated DASIT^EGFP-PURO^ into murine *Bcr-Abl*-driven mouse acute B-cell lymphoblastic leukemia cells (B-ALL) at a low MOI. These cells were then transduced with a mouse base editing sensor library (MBESv2) that contains sgRNAs designed to engineer cancer-associated mutations.^49^

To establish baseline library representation, cells were sampled for next-generation sequencing (NGS) and subsequently electroporated with an optimized adenine base editor (ABE8e-NG) mRNA on Screen Day 0 (Fig 4A). Editing was allowed to occur over a three-day period, which resulted in 56% of the population activating the EGFP reporter construct (Fig 4B). Cells were then split into an untreated control group or selected with puromycin for 2 days. On Screen Day 5, we collected additional cell pellets to assess gRNA library coverage immediately after selection and to identify early bottlenecking that might be mediated by DASIT selection. We continued the screen until Screen Day 15 and collected the final cell pellets for sequencing. Throughout the screen, we maintained consistent cell counts to preserve at least 1000x library coverage.

**Figure 4.**
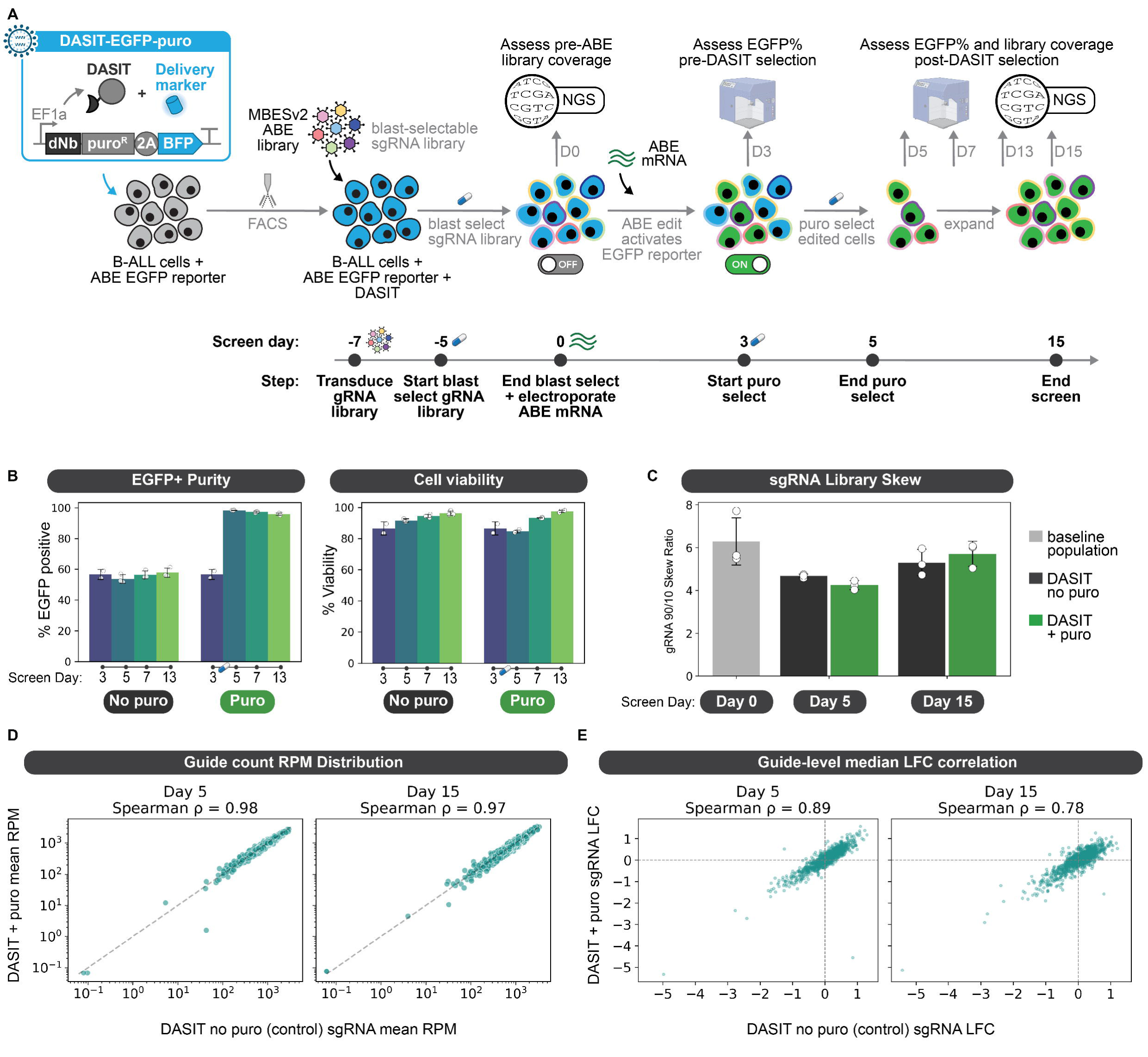
DASIT enables scalable activity-based selection for high-throughput base editing screens. A. Schematic overview of DASIT^EGFP-PURO^ selection for scalable base editing screen setup. B. Percentage of EGFP-positive cells in the no puro control condition vs. the 5 µg/mL puromycin selection condition as assessed by flow cytometry on Screen Days 3, 5, 7, and 13 (*left*). Cells were treated with puromycin for 48 hours between Screen Day 3 and Day 5. Cell viability was assessed with Zombie NIR viability dye (*right*). Data are mean ± s.d. of n = 3 biological replicates. C. Library 90/10 skew ratio (90th percentile guide counts / 10th percentile guide counts). D. Spearman correlation between the gRNA reads per million (RPM) for the no puro control and 5 µg/mL puromycin selection condition. The RPM from each biological replicate (n=3) were averaged for this correlation. E. Spearman correlation of the median log2 fold change for each gRNA from the three biological replicates across the no puro control and 5 µg/mL puromycin selection conditions.

Immediately after the two-day puromycin treatment, we observed robust enrichment for EGFP-positive cells without any subsequent loss of cell viability, and both metrics remained stable throughout the screen (Fig 4B). Sequencing analysis indicated there was no significant skew in sgRNA library representation in cell populations from the control and puromycin conditions at either Screen Day 5 or 15 (Fig 4C). We observed near-perfect correlations in the reads-per-million counts for each sgRNA across the two conditions at both time points (Fig 4D). To compare cellular fitness phenotypes, we computed the median log fold-change values of sgRNAs at different timepoints relative to the input at Screen Day 0. Notably, the strong correlation between control and DASIT-selected libraries consistently held even when comparing cellular fitness phenotypes (Fig 4E). Together, these results show that DASIT enables robust and scalable enrichment of cells with active genome editing while preserving sgRNA library representation and screen outcomes. This approach using DASIT offers substantial time and cost savings over FACS for *in vitro* screens and provides a scalable strategy to select specific cell populations for *in vivo* implantation. Because DASIT-mediated enrichment preserves library representation while eliminating prolonged sorting, this approach enables high-complexity functional screens to be performed at scales that are impractical with traditional FACS-based enrichment. This scalability expands the feasible design space for pooled genome engineering experiments.

### DASIT selects for engineered neurons for acute grafting following spinal cord injury

Cell-fate reprogramming of somatic primary cells often generates mixed cultures of different cell types^51–55^. Isolation of fully reprogrammed cells is essential for phenotype-based applications in drug screening, disease modeling, and cell therapies. We hypothesized that DASIT could enable scalable selection of successfully reprogrammed cells from mixed cultures in cell-fate reprogramming of somatic primary cells. To demonstrate DASIT selection after cell-fate reprogramming of somatic primary cells, we used a highly efficient direct conversion system to generate motor neurons^51,52,56^. To drive conversion, we deliver a virus encoding three motor neuron-specific transcription factors and a high-efficiency chemo-genetic module. After two weeks, cells that successfully convert from Hb9::EGFP transgenic mouse embryonic fibroblasts (MEFs) into induced motor neurons (iMNs) express EGFP at high levels. iMNs are functional and engraftable *in vivo* ^51,52,56^.

Previously, we isolated Hb9::EGFP-positive iMNs using FACS. As FACS provides limited scalability and induces significant cell stress, we hypothesized that DASIT^EGFP-PURO^ could enrich Hb9::EGFP-positive iMNs (Fig 5A-B). At 14 dpi, we measured iMN conversion, quantifying both the total count of Hb9::EGFP-positive cells and the purity of Hb9::EGFP-positive cells. Purity is defined as the total count of Hb9::EGFP-positive cells divided by the total count of cells detected by flow cytometry^22,23^ (Fig 5C). In our initial tests in 96-well format, we observed that selection failed when cells were highly confluent. To prevent overcrowding, we replated cells at 7 days post infection (dpi) at a 1:2 ratio into new wells containing puromycin (Fig S9A). We compared selection with DASIT^EGFP-PURO^ to the wtNb^EGFP^ control and a no-nanobody control. As expected, delivery of DASIT, but not the controls, increases iMN purity in a puromycin dose-dependent manner (Fig S9A). Puromycin treatment reduces the total iMN count (Fig S9A). Potentially, the timing of selection may reduce the total number of iMNs by eliminating cells in the process of conversion which have yet to turn on the reporter or that express EGFP at lower levels^51,53^.

**Figure 5.**
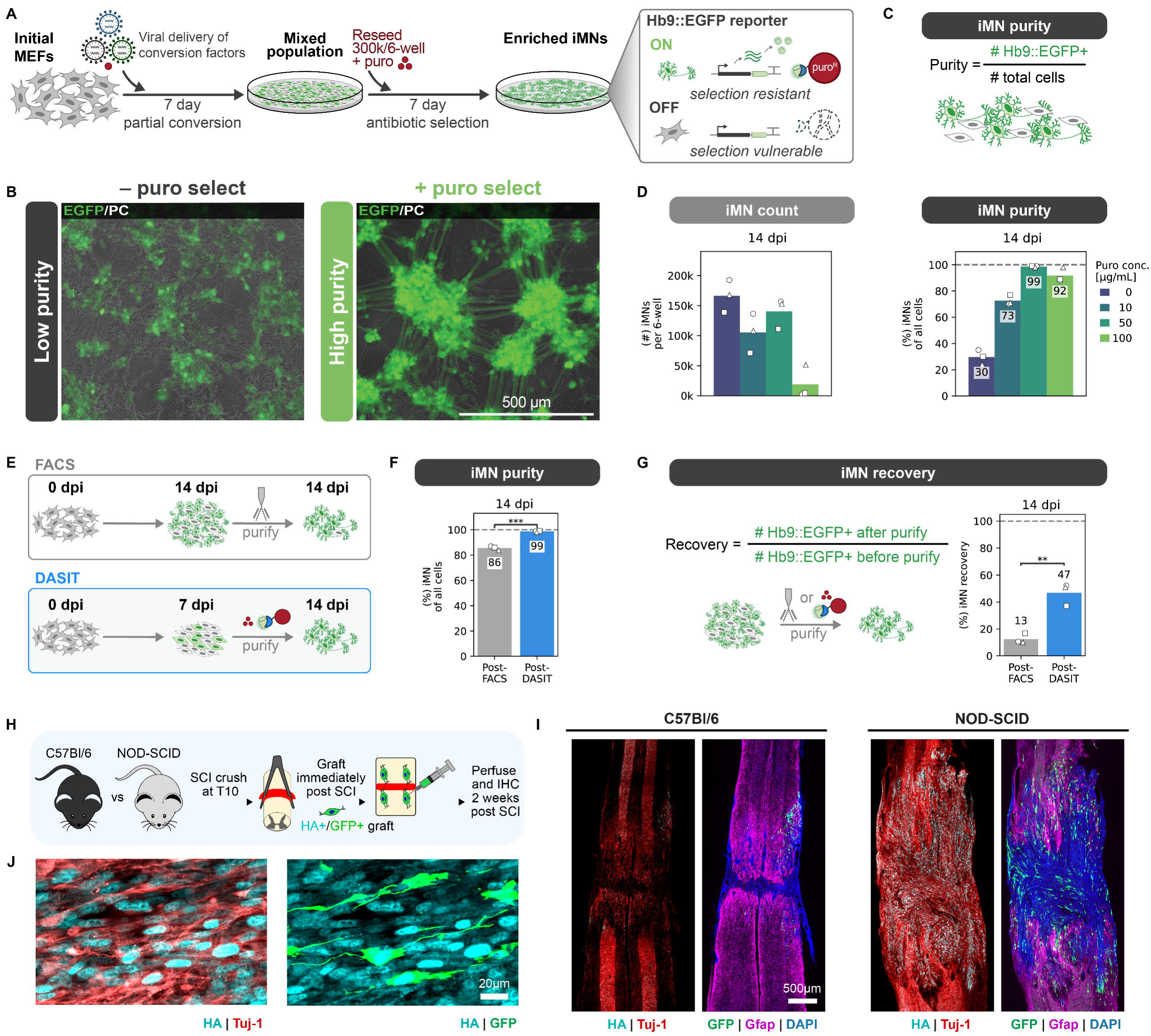
DASIT selects for engineered neurons for acute grafting following spinal cord injury. A. Schematic of reprogramming and selection. One virus encodes three motor neuron-specific transcription factors, a second virus encodes DASIT^EGFP-PURO^, and a third virus and small molecule comprise a high-efficiency chemo-genetic module that increases conversion rates^51,52,56^. DASIT converts EGFP expression from a cell state EGFP reporter (Hb9::EGFP) into a cell state-specific antibiotic resistance. 7 days after direct conversion of MEFs to iMNs is initiated, the mixed Hb9::EGFP-negative and positive culture is reseeded at 300k/6-well Hb9::EGFP+ cells at 7 dpi. B. Images of Hb9::EGFP fluorescence overlaid with phase contrast after 7 days of puro selection with DASIT. C. We define purity as the number of Hb9::GFP+ cells out of all cells detected by flow cytometry and count as the total number of Hb9::EGFP-positive cells per 6-well. D. Conversion outcomes can be quantified by count (*left*) and purity (*right*) after 7 days of puro selection with DASIT^EGFP-PURO^ at different concentrations. Mean is shown and markers denote biological replicates; n = 3 biological replicates per condition. E-G. (E) Schematic of purification by FACS (*top*) vs. DASIT (*bottom*). At 14 dpi, cells were collected to assess (F) purity and (G) recovery. We define iMN recovery as the number of Hb9::EGFP cells that are detected by flow before and after either FACS or DASIT purification. Mean is shown; markers denote biological replicates; n = 3 biological replicates per condition; two-tailed t-test. H. Schematic of acute transplantation of iMNs following SCI crush. NOD-SCID (NOD.Cg-Prkdc^scid^) and C57Bl/6 mice underwent a laminectomy at T10. Acute transplantation was performed immediately post-injury. I. Immunostaining of HA (Ngn2^x3HA^), neuronal marker Tuj-1, and the glial marker Gfap. DAPI labeling of nuclei and HA indicates that grafts survive robustly in NOD-SCID lesions but minimally at C57Bl/6 lesions. Grafts maintain neurogenic potential and extend axons along preserved white matter tracts and longitudinally oriented astrocytes. J. Staining of section of NOD-SCID grafts for HA (Ngn2^x3HA^) and Tuj-1. Significance summary: p > 0.05 (ns), *p ≤ 0.05, **p ≤ 0.01, ***p ≤ 0.001, and ****p ≤ 0.0001.

To increase the iMN numbers for transplantation studies, we scaled up the conversion method to a 6-well format and reseeded a standard count of Hb9::EGFP+ cells per 6-well at 7 dpi, when we started puromycin treatment (Fig 5A, S9B). Titrating the puromycin concentration, we identified a concentration that optimized iMN counts and purity (Fig 5D). We compared iMN purification by FACS at 14 dpi and DASIT selection from 7 to 14 dpi (Fig 5E, S9C). Compared to FACS, DASIT achieves higher purity of iMNs (Fig 5F, S9D). A standard sorting mode was used for FACS to maximize cell yield. Potentially, greater purity can be achieved by sorting with greater stringency but at the cost of reducing total recovery of iMNs. To quantify recovery, we divided the number of Hb9::EGFP-positive cells after purification by the number of Hb9::EGFP-positive cells prior to selection or sorting (Fig 5G). At 14 dpi, we observed a nearly 4-fold increase in iMN recovery of DASIT over FACS (Fig 5G, S9C-E). To identify differences in FACS and DASIT isolated cells, we compared the transcriptional profiles of iMNs purified by FACS and DASIT at 14 dpi using bulk RNA-seq (Fig S10). Normalized gene expression between FACS and DASIT isolated cells is highly correlated (Fig S10A). Gene ontology analysis indicates that DASIT-purified iMNs are enriched for pathways associated with oxidative phosphorylation^57,58^, whereas FACS-purified iMNs enriched genes involved in cell cycle (Fig S10C). Greater purity of DASIT-isolated iMNs may account for the small but significant differences in transcriptional profiles between the bulk transcriptional profiles.

Previously, we showed that *in vitro*-engineered iMNs can graft into the striatum of immunocompetent C57BL/6 mice under both healthy and L-NIO stroke injured conditions^52^. FACS isolation at this scale (>10^6^ iMNs) requires multiple hours of sorting and reduces cell recovery (Fig 5G, S9D). With higher rates of recovery using DASIT, we wondered if we could graft DASIT-selected iMNs into the spinal cords of mice following an acute spinal cord injury (SCI). Grafting neural replacement therapies within the subacute window offers the greatest opportunity to restore lost neural circuits^59–61^. However, for hemorrhagic SCI lesions, cell survival may be compromised when grafted directly into the lesion site due to cytotoxic stressors derived from extravasated blood components, associated inflammation and immune cell infiltration. Elimination of specific immune responses may preserve grafted cell viability^62,63^. Alternatively, fragile neuronal therapies may not survive grafting following an acute SCI regardless of immune activity.

To examine these hypotheses, we used DASIT to isolate engineered motor neurons at scale for grafting in immunocompetent and immunocompromised mice. By eliminating FACS-related cell stress and losses, we could compare neurons grafted with minimal processing-induced stress. To examine the influence of infiltrating immune cells, we compared immunocompetent and immunocompromised mice (C57BL/6 and NOD-SCID mice, respectively). We grafted iMNs into thin crush SCI immediately following injury induction (Fig 5H). We were able to detect grafted iMNs by Hb9::EGFP. Additionally, we could detect an abundance of cells in the immunocompromised mice that are marked by the HA-tag from Ngn2^x3HA^, one of the transcription factors delivered that initiate iMN direct conversion. In the immunocompetent mice, iMN grafts survive poorly and persist locally in the preserved neural tissue immediately adjacent to the lesion site. In NOD-SCID mice, we observe robust survival of grafted cells within the lesion. The iMN SCI grafts showed widespread expression of Nfm and Tuj-1 and extended axon projections that were oriented along preserved axonal tracts in the white matter. The iMNs showed enhanced neuronal fate commitment when compared to an mESC-derived definitive NPC line grafted at equivalent post injury times, which almost exclusively differentiate into Gfap-positive astrocytes^39^. By contrast, we detected no Gfap-positive cells deriving from grafted iMNs, suggesting they are resilient to the cues from acute SCI lesions that can attenuate neurogenic potential. While all HA-positive grafted cells expressed Tuj1, we noted reduced or silenced expression of Hb9::GFP in many of these cells, potentially indicating that grafting influences neuronal identity. Overall, these results demonstrate that DASIT purified iMNs can be produced at transplantation scale and survive after engraftment into SCI-lesions with the neurons persistence limited primarily by immune-mediated rejection.

## DISCUSSION

Altogether, we demonstrate that DASIT supports cell selection for diverse applications. Previous work using desNbs primarily focused on identification of cells and protein interactions, as well as FACS-based isolation^17,32,64–66^. Here, we show that DASIT functions across multiple resistance markers to support scalable enrichment of diverse mammalian cells including human iPSCs, cancer cell lines, and engineered neurons. We show that transient expression of DASIT can be used to generate dual landing pad iPSC lines from a novel iPSC donor, to enrich for cells based on editing activity, and to purify engineered neurons for neuron replacement therapy in spinal cord injury. Overall, DASIT offers a flexible, robust system for cell enrichment that avoids FACS and stable integration of resistance markers, potentially porting into myriad existing workflows to accelerate cell selection.

Direct selection of transgenes via DASIT will fundamentally change how vectors, integration systems, and libraries are designed. In many workflows, DASIT eliminates the need for addition of selection markers which are often included through a self-cleaving tag, internal ribosome entry site or second transcriptional unit^67,68^. Each of these additions increases the size of the cargo which can be a limiting feature for delivery. Additionally, multicistronic elements can alter RNA processing and translation, potentially influencing expression levels and interfering with transgene-associated regulatory processes^67,68^. Expression of selection markers from a separate promoter avoids these confounders, but introduces the potential for undesirable coupling between transcriptional units^69,70^. DASIT selection eliminates these design limits by directly selecting for cells expressing the transgene.

Protein-based circuits like DASIT offer flexibility in delivery without risk of genome integration and avoid the limitations of DNA-based systems. Unlike transcriptional reporters, DASIT directly couples antigen binding to protein stabilization without transcriptional amplification. While traditional selection methods require permanent integration of the selection marker with the synthetic circuit, DASIT antigens can serve as a live reporter of circuit expression for which transient DASIT expression can facilitate enrichment. We show that DASIT supports logic-based installation of landing pads and retains the ability to recycle the selection markers for downstream integration of cargoes in the same cell line.

DASIT offers a scalable method of isolating desired cell phenotypes and fates. We used DASIT to reveal that immune-mediated killing influences neuron survival in grafts delivered within the subacute window. These findings demonstrate that DASIT-selected neurons can survive within the timeframe when grafts are expected to be most effective. Our findings identify the acute immune response as a critical mediator of grafting outcomes, motivating strategies to engineer grafted neurons to mitigate or evade immune detection^62,71–73^. Given the abundance of EGFP-based (and structurally similar fluorescent proteins such as EBFP) reporter systems including transgenic reporters and endogenously tagged proteins, DASIT can be rapidly integrated into many existing workflows for phenotype-, identity-, or state-based enrichment without need for FACS. We show that DASIT supports robust functional enrichment of cells with base-editing activity from heterogeneous pooled cell populations without skewing library representation. By increasing scalability, DASIT opens the potential for larger functional screens *in vitro* and *in vivo*.

Beyond simplifying enrichment, DASIT creates a framework for full automation of cell line engineering. Because DASIT can operate through transient modRNA delivery and antibiotic selection, a minimal set of resistance markers can be rapidly recycled across multiple steps in a workflow. For iterative genome writing workflows^13–16^, DASIT can speed up marker recycling and potentially support multi-step integrations in a continuous, automated workflow. To accelerate transgene integration into landing pads, we envision that the entire process from landing pad installation to transgene integration can be consolidated into a programmable, sequential workflow performed entirely in bulk culture using liquid handling robotics and scalable cell manufacturing platforms. In this configuration, landing pad installation, DNA cargo integration, and excision of auxiliary elements become modular steps within a closed, selection-driven circuit, removing manual bottlenecks and reducing the time to fully engineered, homogenous polyclonal lines from months to weeks. By eliminating FACS dependence and expanding selectable logic beyond conventional antibiotic resistance markers, DASIT transforms DNA integration platforms from a powerful but labor-intensive strategy into a potentially automatable, high-throughput genome engineering system suitable for large-scale screening and programmable cell engineering applications. With appropriate engineering, DASIT selection may also be able to directly select for integrated payloads without an intermediate landing pad installation step. Pairing automation with pooled libraries will massively expand the pace of synthetic biology design-build-learn-test workflows^74^. We envision that this novel approach will facilitate the rapid generation of patient-specific landing pad iPSCs for drug screening and disease modeling^36,75–79^.

In theory, DASIT can be integrated with existing protein-based circuits to expand the types of computations used for selection. Existing protein-based circuits primarily use proteases, degrons, and synthetic peptides to compute responses^19–23,25,80^. While these systems offer broad synthetic expansibility, DASIT’s direct mechanism of sensing protein levels offers a potential mechanism for sensing both synthetic and native protein signals^81^. Protein-based circuits that use multiple inputs of protein levels may more precisely define cell state and fate and can readily integrate with traditional transcriptional reporters. However, detection without compromising protein activity requires that nanobody-antigen interactions do not interfere with protein functions through occlusion of interaction sites and/or mislocalization^82–84^.

As the number of nanobodies expands^27,85^, we envision that DASIT could be extended to new antigens. In particular, selection based on expression of endogenous genes may support enrichment of defined cell types. As we show that the small epitope tag ALFA works effectively with DASIT selection, we envision that cells may be selected based on expression of endogenous genes edited to include ALFA or similar, small tags with cognate antibodies. However, while DASIT offers a programmable and modular framework, we anticipate that there may be challenges for adapting DASIT applications. As selection requires a fine balance between antigen expression and DASIT expression, we expect that some genes may require tuning, and some may not be amenable to DASIT selection. For genes expressing target antigens at lower levels, effective enrichment may require tuning the expression of the desNb to prevent antigen-independent resistance. Subcellular localization of antigens may also influence the efficacy of DASIT selection. Putatively, antibiotic resistance markers localize and function within the cytoplasm to restore protein synthesis or prevent DNA damaging agents from reaching the nucleus^86,87^. Thus, non-cytoplasmic antigens may not effectively stabilize desNbs in the current DASIT architecture. Furthermore, development of alternative selection genes beyond antibiotic resistance markers could expand the repertoire of DASIT for selection of desired cells. Finally, we observed that fusion of destabilized nanobodies to resistance markers requires empirical testing to validate effective DASIT designs. Directed evolution techniques may support rapid identification of novel DASIT fusions^88^.

Overall, DASIT establishes a generalized framework for converting intracellular antigens into selectable markers, paving the way for scalable, flow-free enrichment of diverse cell types across a range of applications. By enabling transient, recyclable selection that operates at the protein level, DASIT reduces the genetic footprint of engineered cell lines and simplifies vector design constraints, features that are particularly advantageous for therapeutic cell manufacturing and genome engineering pipelines.

## ACKNOWLEDGEMENTS

Research reported in this manuscript was supported by the National Institute of General Medical Sciences of the National Institutes of Health under award number R35GM143033 to K.E.G. and R35GM154942 to T.M.O.. Additional funding from the National Science Foundation under the NSF-CAREER under award number 2339986, and by the Air Force Research Laboratory MURI (FA9550-22-1-0316) and the Pershing Square Foundation. N.B.W. and M.E.E. are supported by the National Science Foundation Graduate Research Fellowship Program under grant No. 1745302. The authors acknowledge the Luria HPC resources that have contributed to the research results reported within this paper. We thank the Koch Institute’s Robert A. Swanson (1969) Biotechnology Center (National Cancer Institute Grant P30-CA14051) for technical support, specifically the Flow Cytometry Core Facility. Work in the Sánchez-Rivera laboratory is supported by the Howard Hughes Medical Institute (Hanna Gray Fellowship, GT15656), the National Cancer Institute (Cancer Center Support Grant P30-CA014051, NCI 1P01CA291694–01A1), and the Koch Institute Frontier Research Program. H.C. is supported by NIH T32GM136540. We thank Kasey Love and Christopher Johnstone for feedback on the development of the manuscript. We thank Zahmiria Johnson, Amy Oh, and Anya Lee for discussions and cloning support in the development of DASIT. We thank Jonuelle Acosta, Manuel Contreras, and Michael Hemann for scientific discussions and conceptual support.

## AUTHOR CONTRIBUTIONS

N.B.W. designed, performed and analyzed data for the experiments for all nanobody constructs in 293Ts. N.B.W. designed the modRNA DASIT versions and assisted with testing and analyzing data for iPSC LPs. N.B.W. designed, performed, and analyzed data for the mouse motor neuron conversion and prepared motor neuron cell grafts. A.B.A designed, performed and analyzed the data from the experiments for generating human iPSC lines containing a landing pad using DASIT. D.S.P. contributed to conceptualization and molecular cloning of DASIT epitopes. D.R.G. contributed to the characterization of DASIT delivery methods and LP line development. M.E.E. contributed to the characterization of DASIT stabilization with diverse nanobodies. M.F.C. contributed to sequencing analysis of iMNs. T.M.O. performed the striatum injections and analysis of cell grafting. N.B.W. and K.E.G. wrote the manuscript. H.C. performed and analyzed base editing experiments with supervision from F.J.S.R.. K.E.G. supervised the project.

## DECLARATION OF INTERESTS

A patent related to this technology has been filed by the Massachusetts Institute of Technology.

### LEAD CONTACT AND MATERIALS AVAILABILITY

#### Lead contact

Requests for further information and resources should be directed to and will be fulfilled by the lead contact, Kate E. Galloway, PhD.

#### Materials availability

Plasmids generated in this study will be available from Addgene.

#### Data and code availability

Any additional information required to reanalyze the data reported in this paper is available from the lead contact upon request.

