## Supplement for "Programmable nanobody circuits for cell selection"

### SUPPLEMENTARY FIGURES

#### Supplementary figure 1.

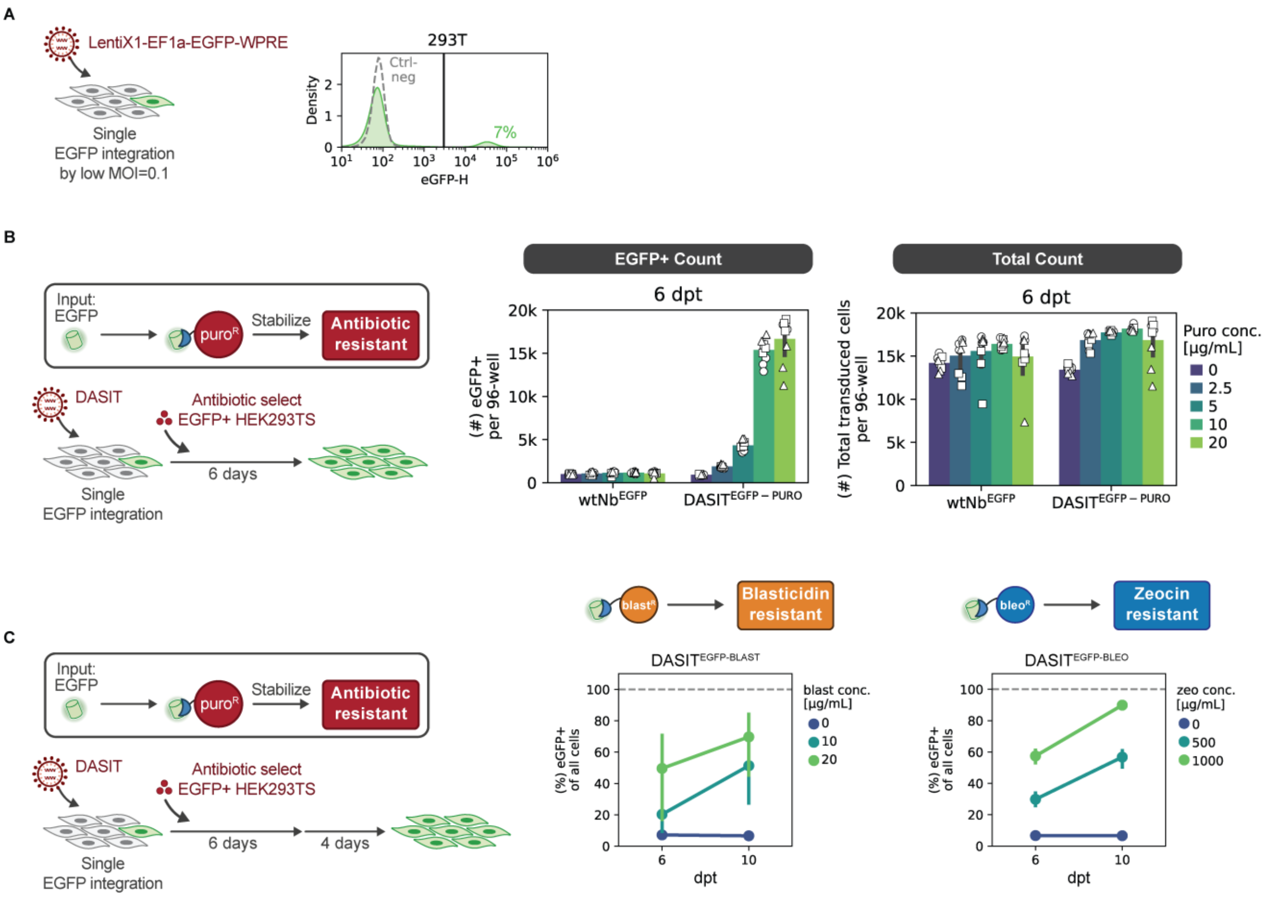


**Supplementary figure 1. Intermediate levels of DASIT expression support DASIT selectivity.**

A. Single EGFP integration in HEK293T was achieved by low MOI=0.1. ~7% of HEK293Ts are EGFP-positive by flow cytometry after expansion.
B. Count data of (*left*) the EGFP-positive cells of transduced cells and (*right*) all transduced cells per 96-well from Fig 1D after 6 days of puromycin treatment across different concentrations. Mean is shown and markers denote biological replicates; n ≥ 3 biological replicates per condition.
C. EGFP-positive purity per 96-well from Fig 1E after 6 and 10 days of antibiotic treatment across different concentrations for (*left*) blasticidin and (*right*) zeocin. n = 3 biological replicates per condition; error bar denotes 95% confidence interval.

### Supplementary figure 2.

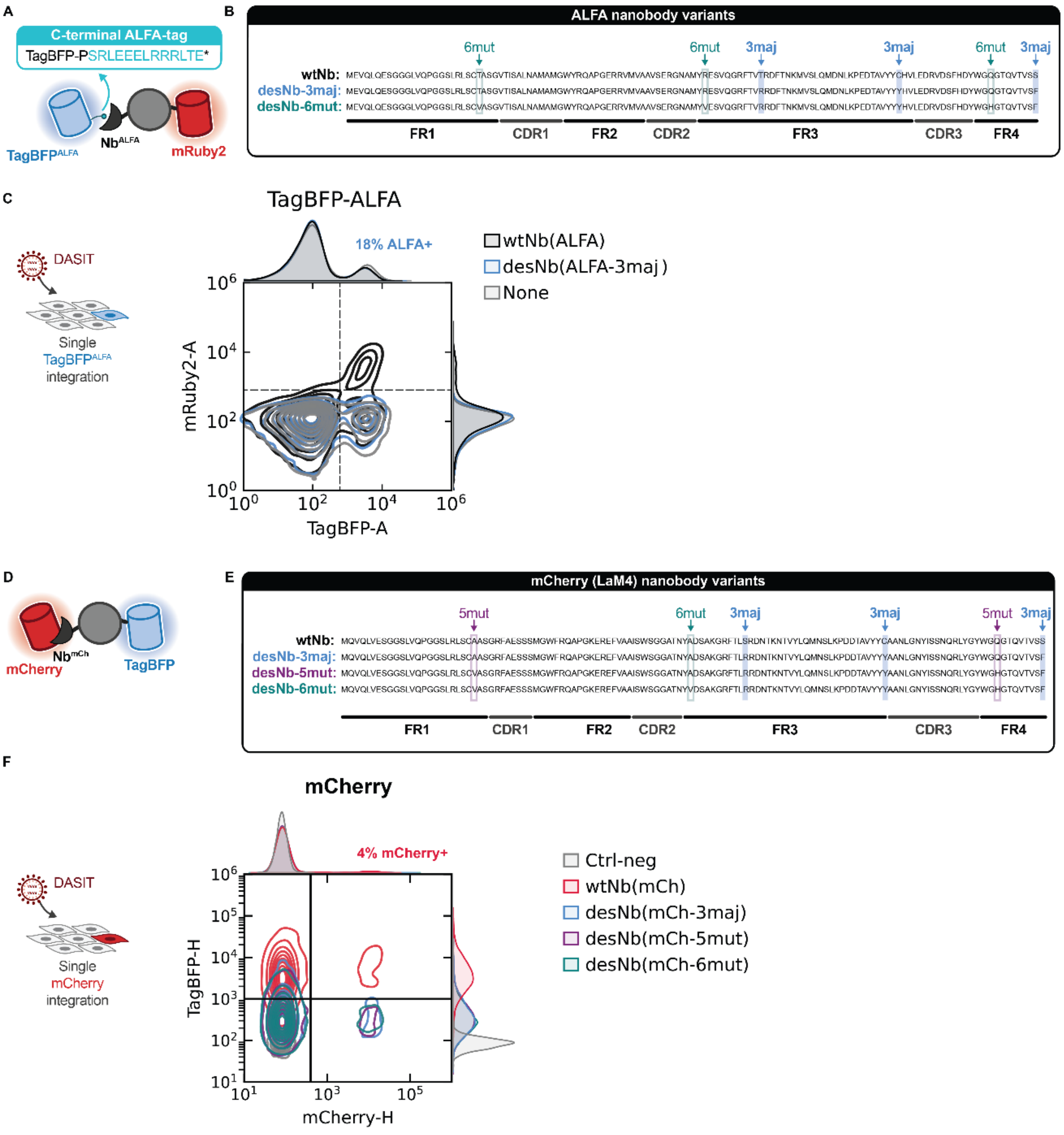


**Supplementary figure 2. DASIT can be adapted to new nanobodies but requires empirical testing.**

A-B. (A) Schematic for detection of ALFA-tag-specific DASIT nanobody stabilization using a C-terminal fusion of ALFA-tag (13 residues: SRLEEELRRRLTE) to TagBFP using mRuby2 fluorescence. (B) Destabilization mutations introduced into the framework regions (FR) of nanobodies should enable rapid generation of destabilized nanobodies from preexisting nanobodies. Three major destabilizing mutations can be used (3maj), however, three additional mutations (6mut) may destabilize the nanobody further.

C. Contour plot of mRuby2 vs. TagBFP fluorescence in HEK293T transduced cells with $TagBFP^{ALFA}$ and different nanobody fusions with either: (black) wtNb(ALFA), (blue) desNb(ALFA-3mut) or (grey) a none (i.e. untransduced) control.

D-E. (D) Schematic for detection of mCherry-specific DASIT nanobody stabilization using TagBFP fluorescence. (E) Different numbers of destabilization mutations were introduced into the framework regions (FR) of the wt LaM4 nanobody to generate destabilized nanobodies targeting mCherry. Three major destabilizing mutations were used for desNb(mCh-3maj), and two to three additional mutations were introduced to destabilize the nanobody further for desNb(mCh-5mut) and for desNb(mCh-6mut), respectively.

F. Contour plot of TagBFP vs. mCherry fluorescence in HEK293T transduced cells with mCherry and different nanobody fusions with either: (red) wtNb(mCh), (blue) desNb(mCh-3maj), (purple) desNb(mCh-5mut), (teal) desNb(mCh-6mut), or a negative control.

### Supplementary figure 3.

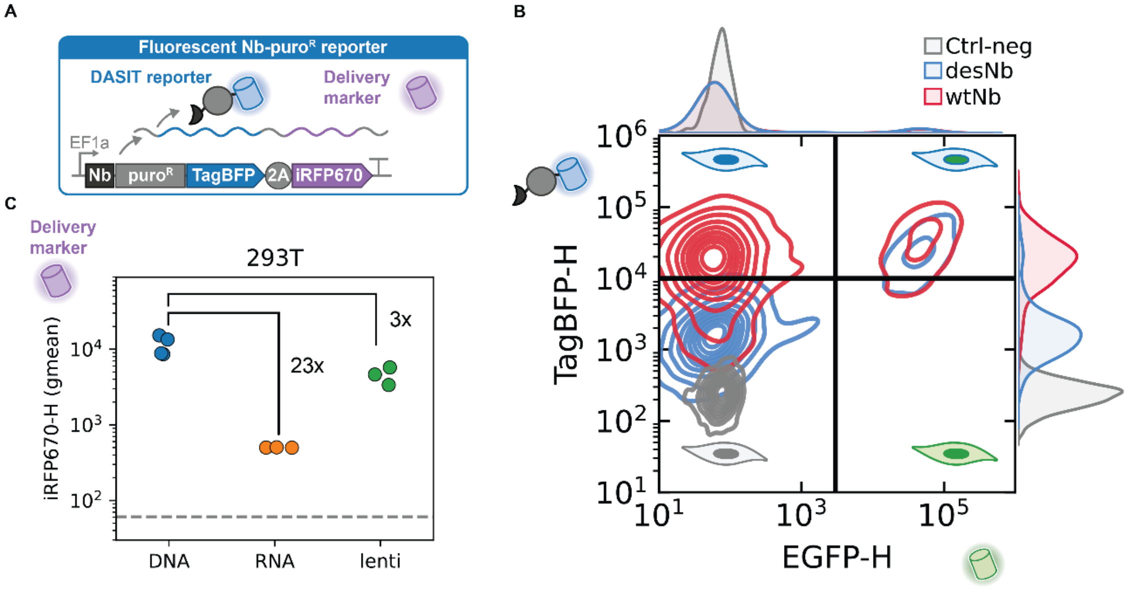


**Supplementary figure 3. Intermediate levels of DASIT expression support selection across stable and transient delivery modalities.**

A. TagBFP fluorescence enables live identification of DASIT expression and a 2A-iRFP670 marker can be used as a co-expression marker.
B. Contour plot of TagBFP vs. EGFP expression measured by flow cytometry lentiviral transduction with DASIT. Ctrl-neg is an untransduced control.
C. DASIT expression differences across modalities. (B) Schematic of DASIT reporter and delivery marker. (C) iRF670 fluorescence distribution. DNA was measured at 48 hpt, RNA at 24 hpt, and lentivirus >6 dpi. (D) Geometric mean for gated iRFP670+ cells across modalities for desNb of iRFP670, the delivery marker. Dashed line indicates the negative control fluorescence. n ≥ 3 biological replicates per condition.

### Supplementary Figure 4.

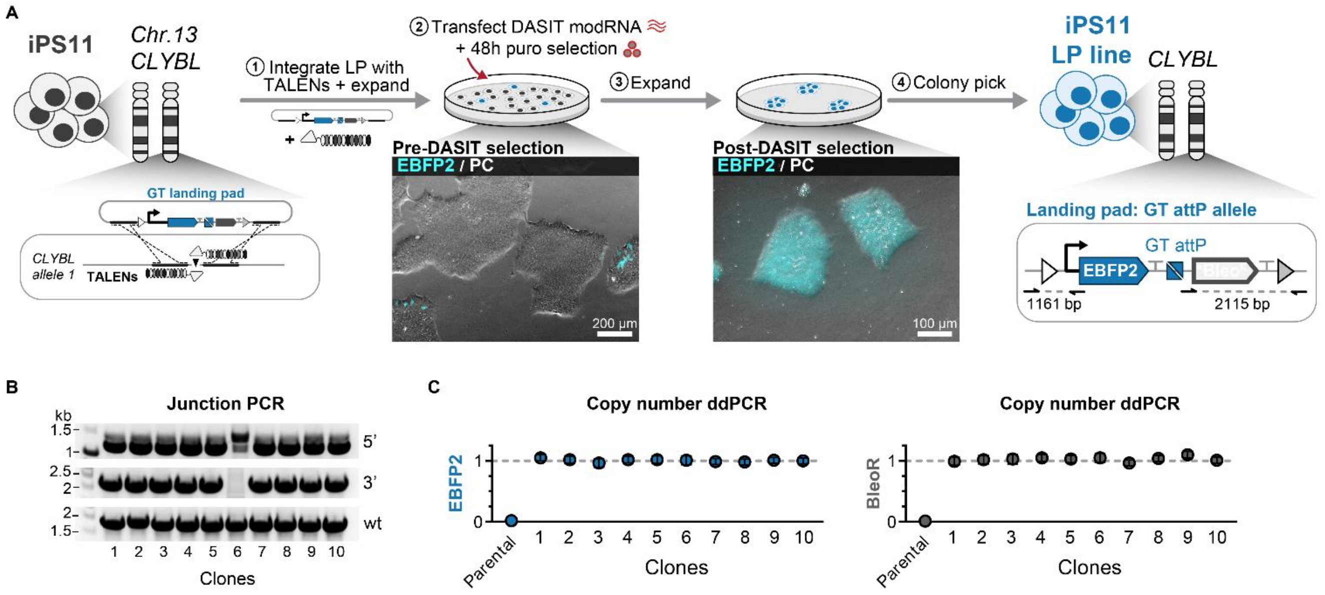


**Supplementary Figure 4. Transient delivery of DASIT for iPSC landing pad creation enables seamless targeting.**

A. Schematic and overlay of fluorescence and phase contrast images before and after DASIT selection procedure. Due to their structural similarities, DASIT$^{EGFP\text{-}PURO}$ can be stabilized by EGFP and its related fluorescent proteins, like EBFP2. As in previous strategies, we used transcription activator-like effector nucleases (TALENs) to generate a double-stranded break at *CLYBL* and integrated the landing pad acceptor DNA (CAG-EBFP2, GT attP, unexpressed *bleoR) by homology-directed repair (HDR) in iPS11. EBFP2 fluorescence in a no TALENS control was used to track dilution of unintegrated landing pad acceptor DNA. After sufficient dilution (~4-10 days), modRNA encoding DASIT$^{EGFP\text{-}PURO}$ was transfected and puromycin selection was started 8 to 24 hours afterward and kept until a no transfection control completely died (48 hours). Surviving iPSCs were expanded and colony picked after growing to sufficient size (~100 µm).
B. Junction PCR analysis confirming targeting of the GT landing pad in the *CLYBL* locus.
C. ddPCR validating single-copy integration in the genome of the GT landing pad. The parental iPS11 cells serve as a negative control for both transgenes. Error bars denote Poisson 95% confidence interval.

**Supplementary Figure 5.**

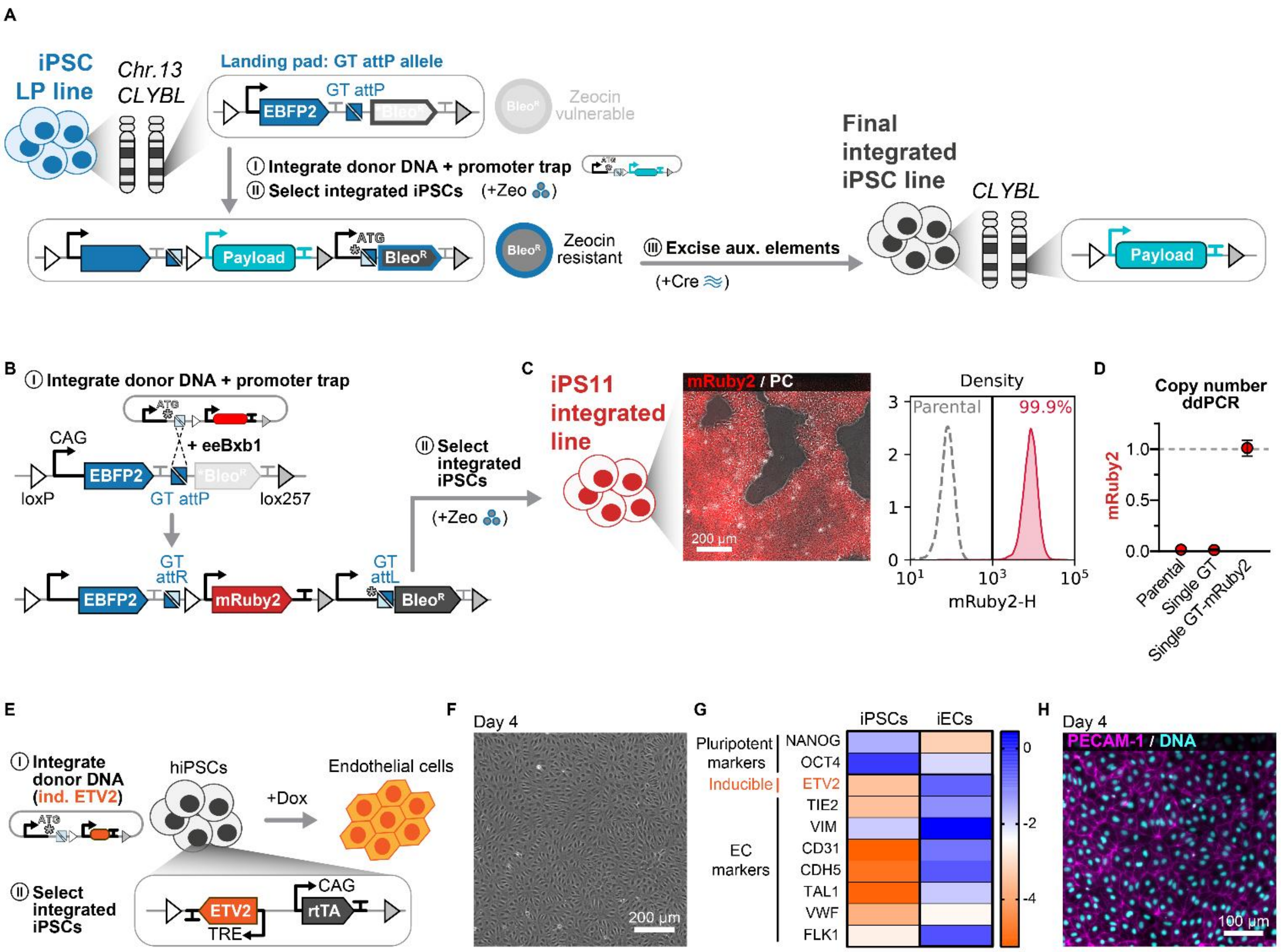


**Supplementary Figure 5. A DASIT-engineered human iPSC landing pad line supports rapid integration of diverse, functional cargoes.**

A. Schematic of STRAIGHT-IN procedure including DNA payload integration and excision of auxiliary elements.
B. Schematic of genomic integration into the GT landing pad (LP) of two DNA payloads, a constitutively expressed mRuby2 fluorescent reporter and a doxycycline-inducible transcription factor ETV2 (ind. ETV2) to drive rapid forward programming into endothelial cells.
C. Overlay of fluorescence and phase contrast image (*left*) and flow cytometry analysis (*right*) showing mRuby2 reporter expression following integration of GT donor plasmid into GT LP. Dashed line denotes untransfected, parental STRAIGHT-IN GT LP iPS11 cells.
D. ddPCR validating single-copy integration in the GT LP of a DNA payload carrying mRuby2 reporter. The parental iPS11 and GT LP cells serve as a negative control for the transgene. Error bars, Poisson 95% CI.
E. Schematic of the engineered ETV2 inducible line established from the DASIT selected STRAIGHT-IN GT LP iPS11 monoclonal line.
F. Representative phase contrast image of iPSCs directed into endothelial cells in the presence of doxycycline for 4 days.
G. Gene expression analysis by qPCR of endothelial markers from the cells in (F) after 4 days of ETV2 induction. Values are normalized to *RPL37A* and shown relative to the uninduced condition ($log_{10}$-transformed). n = 3 independent differentiations.
H. Immunofluorescence images of ETV2 inducible iPS11 line after 4 days of ETV2 induction, cultured with doxycycline (PECAM-1, magenta; DNA, cyan).

### Supplementary figure 6.

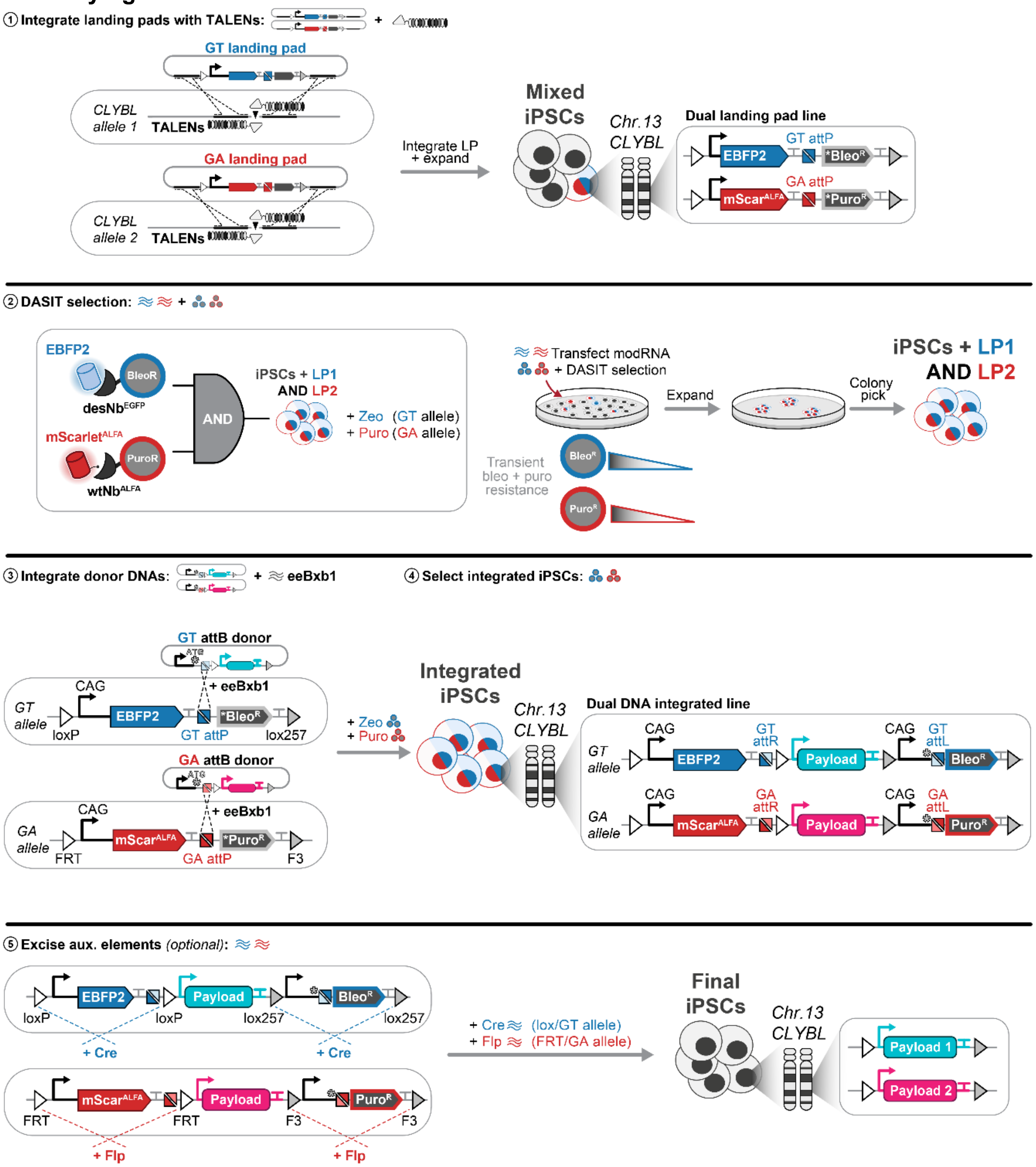


**Supplementary figure 6. Overview of the generation of iPSC landing pad lines using DASIT and STRAIGHT-IN workflow.**

Schematic for generating STRAIGHT-IN DUAL lines in human iPSCs at the *CLYBL* locus using DASIT.

**Step 1: Integrate landing pads using TALENs**. A plasmid encoding TALENs targeting *CLYBL*[47] is transfected into human iPSCs with the two targeting constructs containing homology arms for *CLYBL*. The targeting constructs are distinguished by orthogonal “GT attP” and “GA attP” recombination sites with a fluorescent EBFP2 and mScarlet[ALFA] reporter, respectively. They also contain promoterless bleomycin resistance and puromycin resistance genes, respectively. Cells are expanded to dilute out the transient expression from the targeting constructs and to allow integrated cells to grow.
**Step 2: DASIT selection**. modRNA encoding DASIT[EGFP-BLEO] and DASIT[ALFA-PURO] are delivered to human iPSCs which confers transient zeocin and puromycin resistance to the cells that integrated the “GT attP” and “GA attP” landing pads. After 3 days of zeocin and puromycin selection, only integrated cells remain which form colonies that can be picked for clonal isolation.
**Step 3-4: Integrate DNA payload and select integrated cells**. Donor plasmids containing matching “GT attB” and “GA attB” recombination sites are transfected, along with the evolved serine recombinase, eeBxb1[89]. Using a promoter trap strategy whereby the donor plasmids contain a promoter and the ATG initiation codon for the selection cassettes, only the cells that integrate the donor plasmid will be conferred resistance to zeocin (bleomycin) and puromycin. After 3 days of zeocin and puromycin selection, only integrated **cells** should remain.
**Step 5: Excise auxiliary elements**. Auxiliary elements from the landing pad and donor plasmids can be excised by transfecting modRNA containing Cre and Flp which will recognize the orthogonal loxP/lox257 and FRT/F3 sites on the GT and GA landing pads, respectively. Upon excision, the cells should only contain the two DNA payloads integrated with small flanking recombination sites.

### Supplementary figure 7.

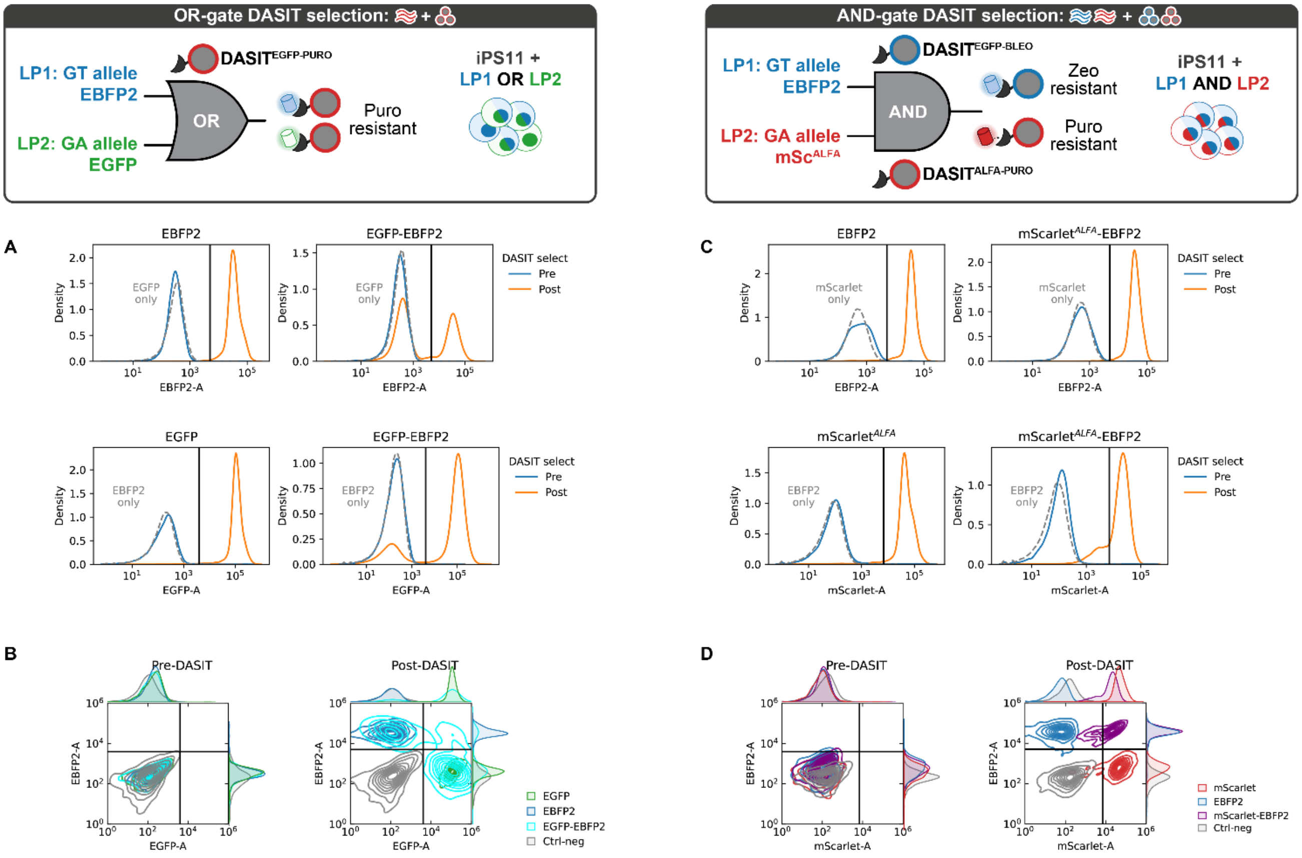


**Supplementary figure 7. Rapid generation of iPSC landing pad lines using DASIT.**

A-B. Representative fluorescent profiles by flow cytometry of EBFP2 (GT allele) and EGFP (GA allele) for the landing pad integration using the OR-gate strategy. The individual distributions are shown in (A) while joint contour plots are shown in (B) before DASIT selection (*left*) and after DASIT selection (*right*) with puromycin.

C-D. Representative fluorescent profiles by flow cytometry of EBFP2 (GT allele) and mScarlet$^{ALFA}$ (GA allele) for the landing pad integration using the AND-gate strategy. The individual distributions are shown in (C) while the joint contour plots are shown in (D) before DASIT selection (*left*) and after DASIT selection (*right*) with puromycin and zeocin.

**Supplementary figure 8.**

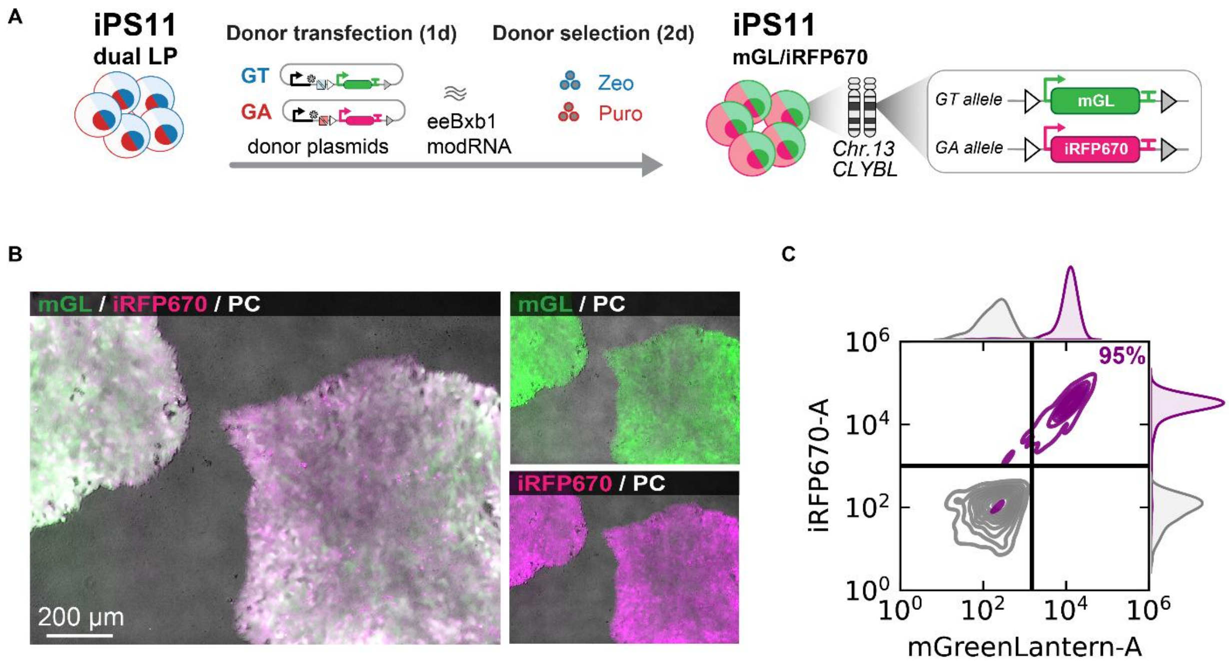


**Supplementary figure 8. Rapid, simultaneous integration of two cargoes in the DASIT-isolated iPS11 dual landing pad lines.**

A. Schematic of STRAIGHT-IN Dual simultaneous integration of two DNA payloads carrying mGreenLantern (GT Donor) and iRFP670 (GA Donor) fluorescent reporters.

B-C. (B) Overlay of fluorescence and phase contrast images and (C) flow cytometry analysis showing mGreenLantern and iRFP670 reporter expression after STRAIGHT-IN Dual payload integration.

### Supplementary figure 9.

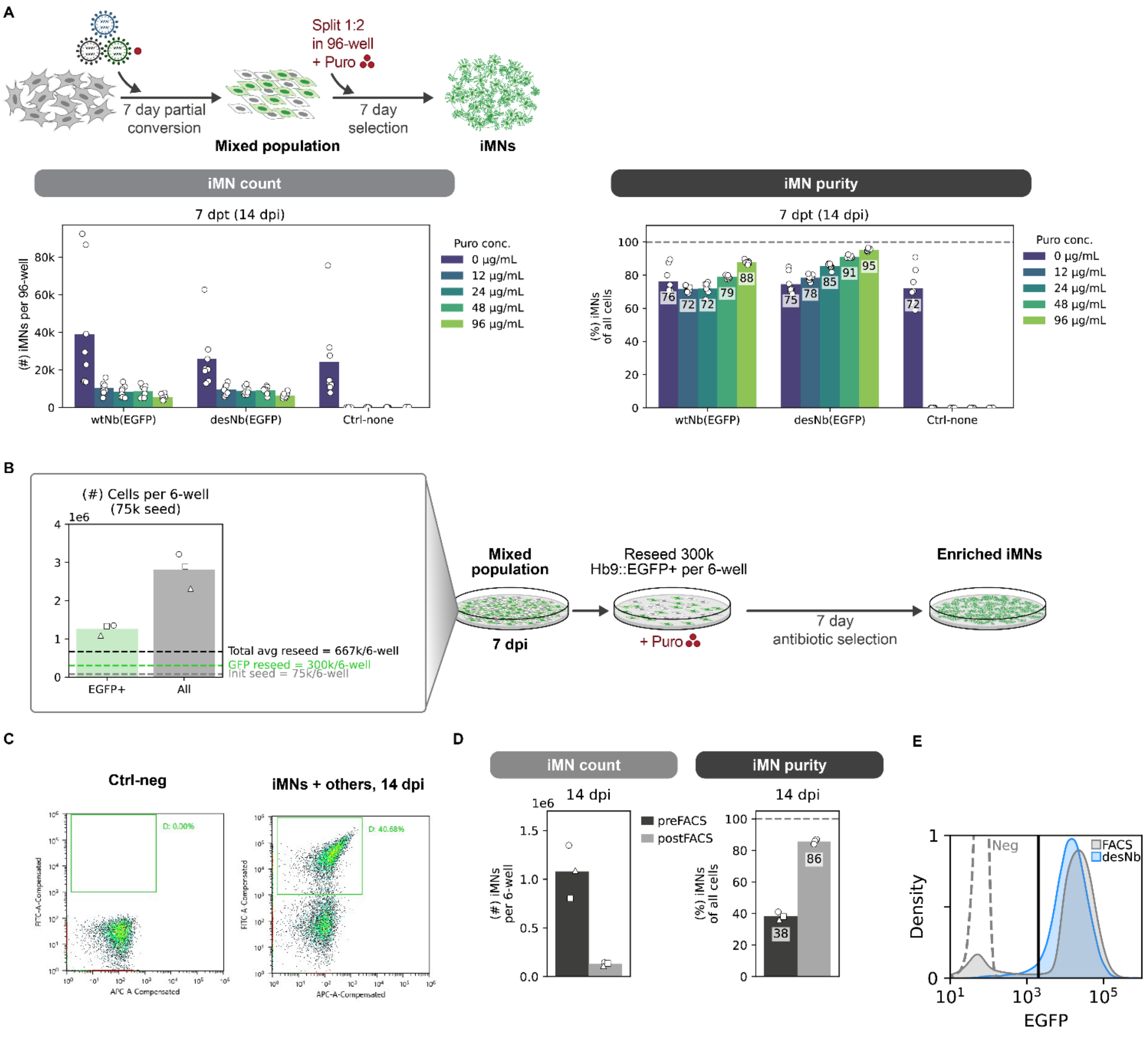


**Supplementary figure 9. DASIT enables selection based on expression of a cell state-specific GFP reporter in primary cells with improved cell recovery compared to FACS.**

A. Schematic of reprogramming and selection at smaller scale in 96-well plates. 7 days after direct conversion of MEFs to iMNs is initiated, the mixed Hb9::EGFP-negative and positive culture was reseeded and split at a 1:2 ratio into new 96-wells until assayed at 14 dpi for quantification. (*left*) total iMN count and (*right*) purity per 96-well at varying puro concentrations for the wtNb(EGFP) control, desNb(EGFP), and an uninfected control. Mean is shown; n = 2-3 biological replicates per condition; dashed line for purity marks 100% purity.

B. Total cell counts on a per 6-well basis at 7 days after conversion is initiated. Dashed lines mark (grey) the initial seeding density of 75k MEFs per 6-well, (green) the reseed density of 300k Hb9::EGFP cells per 6-well at 7 dpi, and (black) the total average reseed density per 6-well at 7 dpi based on a goal of 300k Hb9::EGFP per 6-well and the EGFP+ percentage of each replicate. Mean is shown for the EGFP-positive and total cell count per 6-well; markers denote biological replicates; n = 3 biological replicates per condition.

C. Representative sorting gate for FACS at 14 dpi for (*right*) mixed conversion population without selection and (*left*) negative control. The fluorescence of EGFP (FITC) vs. an unused autofluorescence channel (APC) was used to gate Hb9::EGFP+ iMNs.

D. (*Left*) iMN count per 6-well and (*right*) purity before and after FACS sorting using the gate shown in Fig S9C. Mean is shown and markers denote biological replicates; n = 3 biological replicates per condition; dashed line for purity marks 100% purity.

E. Fluorescence distribution for iMNs after purification by (blue) DASIT$^{EGFP\text{-}PURO}$ and (grey) FACS with a (grey dashed) negative control. Black line denotes the Hb9::EGFP+ thresholding for iMNs.

### Supplementary figure 10.

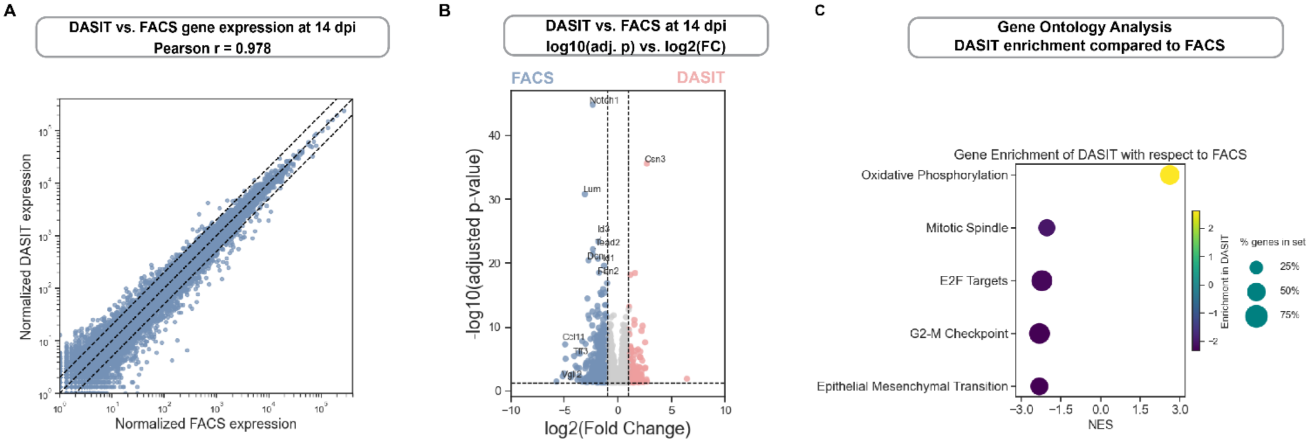


**Supplementary figure 10. Transcriptional profiling of iMNs selected via DASIT and FACS indicates strong correlation of profiles.**

A. Correlation of gene expression between DASIT and FACS-purified iMNs at 14 dpi. Normalization of gene expression calculated via DESeq2 “median of ratios”. Dashed lines indicate x=y with two-fold change on either side. Each point denotes the mean normalized read count across n = 3 biological replicates.
B. Volcano plot from transcriptional profiles between DASIT and FACS-purified iMNs at 14 dpi.
C. Gene ontology analysis showing 5 most differentially enriched pathways in DASIT vs. FACS-purified iMNs at 14 dpi, as indicated by Normalized Enrichment Score (NES). For all pathways, adj. p-value < 0.0001; n = 3 biological replicates.
